# The effect of developmental variation on expression QTLs in a multi parental *C. elegans* population

**DOI:** 10.1101/2023.04.21.537811

**Authors:** Bram van Eijnatten, Mark Sterken, Jan Kammenga, Harm Nijveen, Basten L. Snoek

## Abstract

Regulation of gene expression plays a crucial role in the development and adaptation of organisms to changing environments. A population-based procedure used to investigate the genetic regulation of gene expression is eQTL mapping. Typically, the age of the population used for eQTL mapping at the time of sampling is strictly controlled. This is necessary because the developmental process causes changes in gene expression, complicating the interpretation of eQTL mapping experiments. However, organisms can differ in their “developmental age”, even if they are of the same chronological age. As a result, eQTL patterns are affected by uncontrolled developmental variation in gene expression. The model organism *C. elegans* is particularly suited for studying the effect of developmental variation on eQTL mapping patterns. In a span of days *C. elegans* transitions from embryo through four larval stages to adult while undergoing massive changes to its transcriptome. Here we use *C. elegans* to investigate the effect of developmental age variation on eQTL patterns and an available normalization procedure. We used dynamical eQTL mapping, which includes developmental age as a cofactor, to separate the variation in development from genotypic variation and explain variation in gene expression levels. We compare classical single marker eQTL mapping and dynamical eQTL mapping using RNA-seq data of ∼200 multi-parental recombinant inbred lines of *C. elegans*. The results show that many eQTLs are actually caused by developmental variation, that most trans-bands are associated with developmental age and that dynamical eQTL mapping detects additional eQTLs not found with classical eQTL mapping.

## Introduction

Regulation of gene expression is a key mechanism underlying the huge diversity of phenotypes, adaptations, and developmental stages within and across species. Understanding gene expression regulation therefore provides crucial insights into the way organisms develop and adapt to changing environments^1–7^. The genetic regulation of gene expression can be investigated through expression quantitative trait locus (eQTL) mapping, also called genetical genomics^8^. In this statistical procedure, polymorphic regions (eQTLs) are associated with variation in gene expression, pinpointing potential regulatory regions. The nematode *Caenorhabditis elegans* is often used in eQTL studies and has a small, well annotated genome as well as well documented genetic variation^9–13^. Its tolerance of cryo- preservation, large brood size and self-fertilizing ability allow for the construction of perpetual recombinant inbred lines (RILs) derived from genetically diverse *C. elegans* isolates^11, 13, 14^. RILs are homozygous for almost all loci and constitute a genetic mosaic of the parental genotypes, thereby increasing the resolution and power of methods for eQTL detection^12^. The above qualities make *C. elegans* an exceptional model system for genetical genomics.

As in most organisms, the developmental process in *C. elegans* is associated with massive changes in gene expression^1, 15, 16^. Many genes are expressed in a developmental stage-specific manner^6, 7, 17–19^. Others are up- and down-regulated cyclically as *C. elegans* transitions through the four larval stages^20–22^. The speed of development, as well as the process of aging, varies between individual *C. elegans* strains due to stochastic factors, maternal age and variation in genetic background^1, 2, 18, 19, 23–26^. Due to the impact of the developmental process on gene expression combined with the interaction with the genetic background, eQTL patterns can be affected by uncontrolled developmental variation^1, 2, 18^. To conceptualize developmental variation, it is useful to distinguish between the chronological age and the developmental age. The chronological age is simply the measured age of the organism, whereas the developmental age represents how far the organism has progressed along the developmental process. Because variation in developmental speed and thus developmental age depends partly on genetic factors, it is intrinsic to populations used for genetic mapping. Careful synchronization of worms at a particular developmental checkpoint can help to reduce but does not eliminate developmental age variation^27^. As a result, the eQTL mapping procedure could attribute expression variation resulting from variation in developmental age to a genetic polymorphism. Conversely, developmental age variation could obscure the effects of the genetic background on gene expression.

One way to deal with the confounding effect of the developmental process in eQTL mapping is to apply a normalization procedure^19, 28^. Another approach is to make the effects of developmental variation on gene expression explicit by including the developmental age as a cofactor in the statistical model for eQTL mapping (dynamical eQTL mapping)^2^. The latter approach is more informative because it explicitly models how eQTLs affect the dynamics of gene expression, rather than just the magnitude.

A seminal paper by Francesconi and Lehner^2^ showed that dynamical eQTL mapping using the quantified developmental age can be leveraged to detect additional eQTLs. By relating the magnitude of gene expression to the developmental age the authors showed that eQTLs can affect the expression dynamics over the course of the developmental process.

In this study we aimed to quantify the difference between the classical eQTL mapping approach, which does not explicitly consider developmental variation, and dynamical eQTL mapping. To this end, we used RNA-seq data of ∼200 *C. elegans* multi-parental recombinant inbred lines (mpRIL)^24, 29^. RNA-seq samples were obtained 48 hours after bleaching, ensuring there is no variation in chronological age within the population. At the time of sampling the mpRILs were in the L4 larval stage. We started by quantifying the developmental age directly from the gene expression data with a straightforward approach involving principal component analysis (PCA). Next, we performed eQTL mapping using linear models, both with and without the developmental age included as a cofactor. We compared these models quantitatively, by the number of eQTLs detected and the variance in gene expression attributed to the eQTLs, and qualitatively, by the distribution of eQTLs over the genome. We show that most, but not all, *trans*-bands (regulatory hotspots) result from a shared association with the developmental age between the SNP marker and transcript levels of genes involved in developmental processes. We present evidence that such markers affect gene expression indirectly, by influencing the developmental speed. We also show that comparing the results from models with and without the developmental age can help to find loci linked to the developmental age as well as direct regulators of gene expression.

## Results

### Developmental age estimation by principal component analysis

We quantified the developmental age of the multi-parental recombinant inbred lines (mpRIL) by performing a principal component analysis (PCA) on the normalized gene expression counts. Estimating relative differences in developmental age using a PCA was shown to be effective in a previous study^28^. Principal component (PC) 1 explained ∼48% of the gene expression variance and PC2 explained ∼5% (**Figure 1A**). Under the assumption that the developmental age would be the largest contributor to the variance in gene expression, we investigated whether PC1 is a good proxy for the developmental age of the mpRILs. First, we looked at the expression of the yolk protein *vit-2* gene, since it is part of a cluster of 53 genes (developmental indicator genes) that show a robust linear increase in expression during the L4 stage (cluster 1 in Snoek *et al*., 2014). The expression of vit-2 increases as the projection of the mpRILs on PC1 increases (Pearson correlation ∼ 0.91) (**Figure 1A**). Second, we investigated whether the other developmental indicator genes also show a positive correlation with PC1. We found a strong positive correlation between the projection on PC1 and the mean developmental indicator gene expression (Pearson correlation ∼ 0.97) (**Figure 1B**). The association between the developmental indicator genes and PC1 is further verified by high projections of these genes on PC1 (**Figures S1**). We validated the developmental indicator gene approach by taking a larger set of 2050 genes shown to be monotonically rising in a different study (Hendriks *et al.*, 2014). The mpRILs ordered by their projection on PC1 sort the expression of these genes well (**Figures S2**). Third, we determined the developmental ages of the mpRILs using the RAPToR package^23^ (Pearson correlation RAPToR ages and PC1 ∼0.91). The mpRILs ordered by the RAPToR age estimates provided a sorting that was less consistent with the known monotonically increasing expression of the developmental indicator genes (**Figures S3**) compared to PC1 (**Figures S4**). Hence, we used the projections of the mpRILs on PC1 as our developmental age estimates in the subsequent analysis. We conclude that PC1 is strongly associated with the developmental process and can be used to approximate the developmental age of the mpRILs. Furthermore, the biological replicates of the parental genotypes cluster very closely together on PC1 (**Figures S5**). This suggests that the genetic background of the mpRILs is an important cause of developmental age variation. On the other hand, a heritability analysis on PC1 indicated a narrow sense heritability (h_2_) of ∼0.23. A potential explanation for the disparity between the similar projections of the parental duplicates on PC1 and the relatively low h_2_ of PC1 could be that epistatic interactions are important determinants of the developmental age, whereas the narrow sense heritability considers only additive genetic effects.

**Figure 1:**
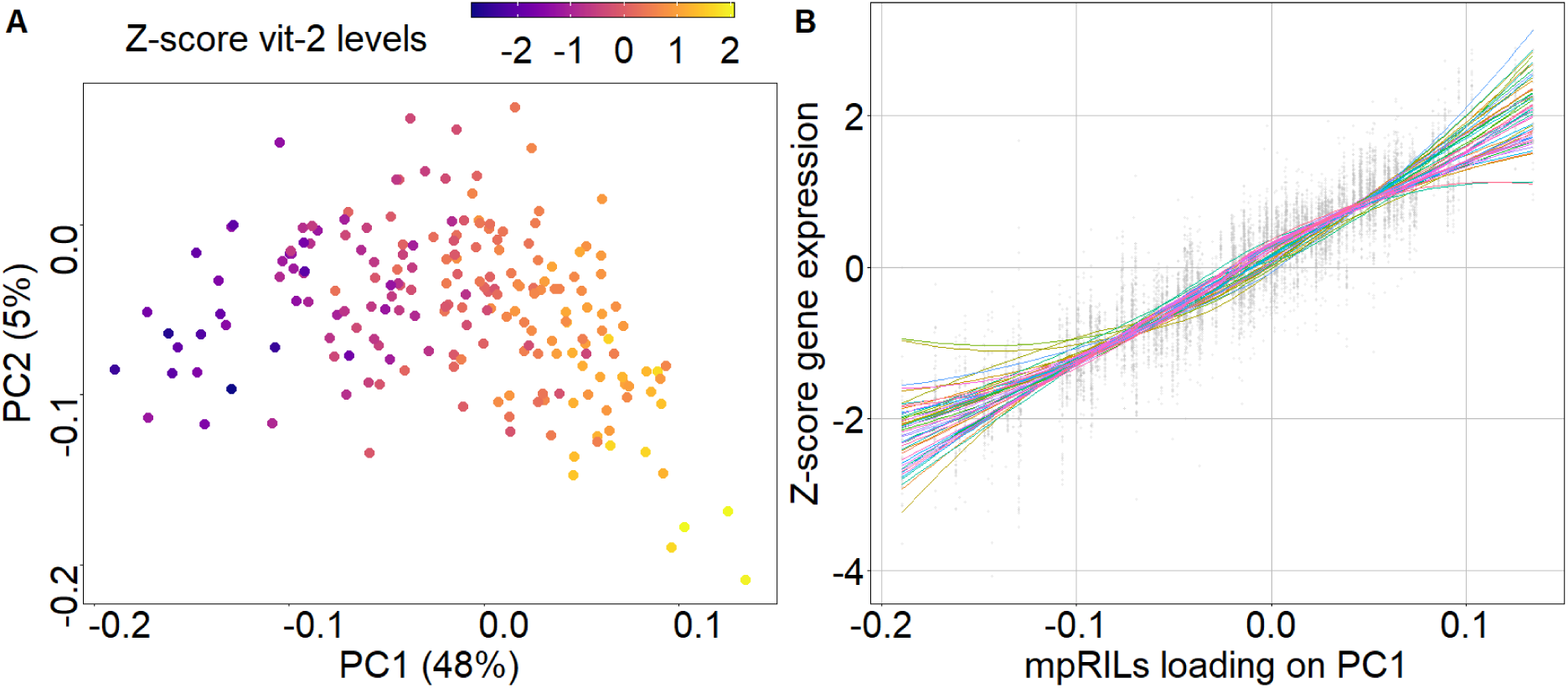
Developmental age estimation using PCA. **A)** PCA results. Points are the projections of the mpRILs on PC1 (x-axis) and PC2 (y-axis). Color indicates the Z-score of the center log ratio of fpkm values of *vit-2*, a developmental indicator gene^1^. **B)** Gene expression of the 53 developmental indicator genes from Snoek *et al.*, 2014 with expression in this dataset. The x-axis shows the projections of the mpRILs on PC1. The y-axis shows Z-score of the center log ratio of fpkm values. Colored trendlines correspond to a generalized additive model fit to the expression values of each developmental indicator gene.

### The effect of developmental age variation on eQTL mapping

To study the effect of variation in developmental age on eQTL mapping experiments we compared a linear single marker model (SMM) with two models including the developmental age as a cofactor, the linear additive age model (AAM) and the linear interaction model (IM). In the case of the interaction model, we called an eQTL if there was a significant marker effect or a significant interaction between the marker and the developmental age. While the interaction model marker (IMM) term detects additive differences in gene expression, a significant interaction model interaction (IMI) term could indicate an eQTL that influences the slope of gene expression over the developmental age. While the models partially overlap, they each also detect a subset of unique eQTLs not detected by the other models (**Figures S6**). We first show the quantitative effect of dynamical eQTL mapping by comparing the SMM with the AAM in terms of number of eQTLs, p-values, strength of the marker effect and the heritability of transcript levels. Next, we show the difference between the models in terms of the distribution of eQTLs over the genome. Finally, we discuss the application of the IM for dynamical eQTL mapping.

Using the AAM, we find 9473 transcripts (out of 12029 polycistronic transcripts) with expression levels affected by the developmental age (-log(p) > 1.79 (FDR = 0.05) (**Table 1**). Finding ∼10,000 transcripts affected by the developmental process is in line with previous reports on the N2 reference strain^1^ and emphasizes the prevalence of developmental variation in gene expression data.

**Table 1:** Significant effects detected by the single marker, additive age and interaction models. An FDR of 0.05 is used unless otherwise specified. The single marker model explains gene expression using the genotype at the marker position. The additive age model explains gene expression using both the marker genotype and the developmental age (PC1) as an additive cofactor. The interaction model explains gene expression using the marker genotype, an additive effect of the developmental age and an interaction between the marker genotype and developmental age. For the interaction model we consider a significant effect of the marker genotype (IMM term) as well as a significant interaction effect (IMI term) an eQTL. * Applies to additive developmental age term of interaction model.

|  | Single marker model (SMM) | Additive age model (AAM) | Interaction model, marker term (IMM) | Interaction model, interaction term (IMI) |
| --- | --- | --- | --- | --- |
| eQTLs detected (FDR <0.05) | 18605 | 8666 | 6886 | 1161 |
| *Cis*/Local | 2136 | 1954 | 1814 | 92 |
| *Trans*/Distant | 16469 | 6712 | 5072 | 1069 |
| *Cis*/*Trans* ratio | 0.13 | 0.29 | 0.36 | 0.08 |
| Transcripts w/ eQTL | 8875 | 6133 | 5180 | 904 |
| eQTLs detected (FDR <0.1) | 27445 | 13369 | 10938 | 4276 |
| Transcripts affected by developmental process | - | 9473 | *8806 | *8806 |

We compared the p-values of the marker effects obtained with the SMM with the p-values of the AAM (**Figure 2A**). We noticed that for a subset of eQTLs (151), the p-value decreases more than 10 orders of magnitude when the developmental age is added to the model (*Strong developmental effect eQTLs*, above top black line in **Figure 2A**). The transcripts with such eQTLs all display a gene expression pattern over the developmental age that has a clear linearly increasing trend and little within allele variation compared to between allele variation (**Figure 3**). Most strong developmental effect eQTLs are detected by both the SMM and AAM. Examining the changes in p-value around the thresholds (**Figure 2B, Table S1**) reveals that many eQTLs are only detected by one of the models. eQTLs that are differentially detected between the models are of special interest as in these cases the qualitative result of the eQTL mapping procedure is impacted, potentially obfuscating or revealing relevant biology. The total number of eQTLs decreases by ∼53% between the SMM (18605) and the AAM (8666) (**Table 1**). The AAM detects 1221 eQTLs that would not be detected by the SMM (*Dynamical eQTLs*) (**Figure 2B cyan color, Table S2**). On the other hand, over 11160 eQTLs (*Single marker model eQTLs*) detected by the SMM appear no longer significantly associated with gene expression when the AAM is applied (**Figure 2B, orange color**). In conclusion, adding the developmental age as a co-factor to the model changes the detection outcome for more than 12000 potential eQTLs.

**Figure 2:**
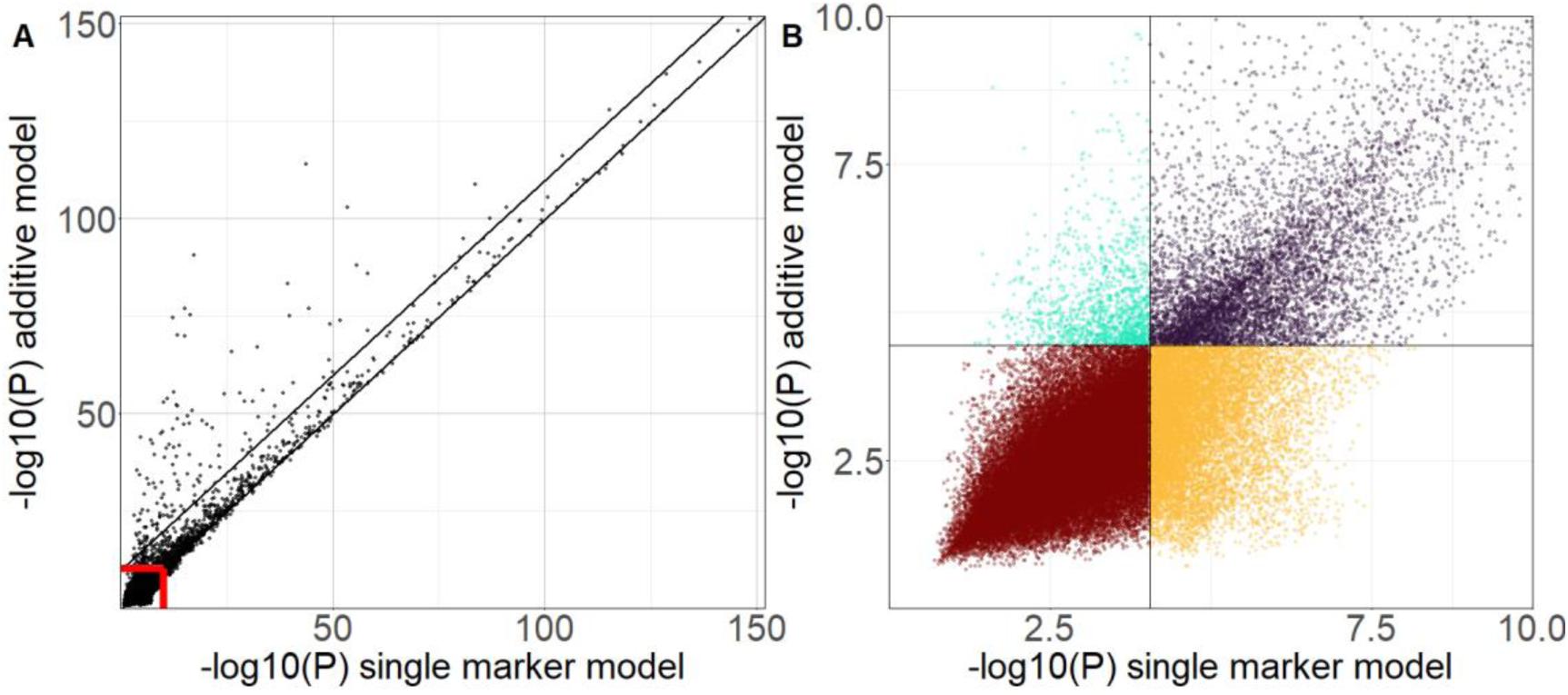
Comparison p-values obtained with the SMM and AAM. **A)** -log10(p-value) of SMM versus AAM, i.e. higher value corresponds to lower p-value. The x-axis shows the lowest marker p-value per chromosome per transcript obtained with the SMM (72174 total plotted p-values). The y-axis shows the same for the AAM. Since we call a maximum of one eQTL per transcript per chromosome, all eQTLs are represented in the plot. The red lines show the subsection of the plot depicted in B. Lower black line is identity line, such that eQTLs above the line have a lower p-value according to the AAM, whereas eQTLs below the line have a lower p-value according to the SMM. EQTLs above the top black line are strong developmental effect eQTLs (P-value ten orders of magnitude more significant with AAM as compared to SMM). **B)** Zoomed-in subsection of A. Points are colored by whether the eQTL was detected with none (brown), both (purple) or one of the models (orange for single marker eQTLs and cyan for dynamical eQTLs).

**Figure 3:**
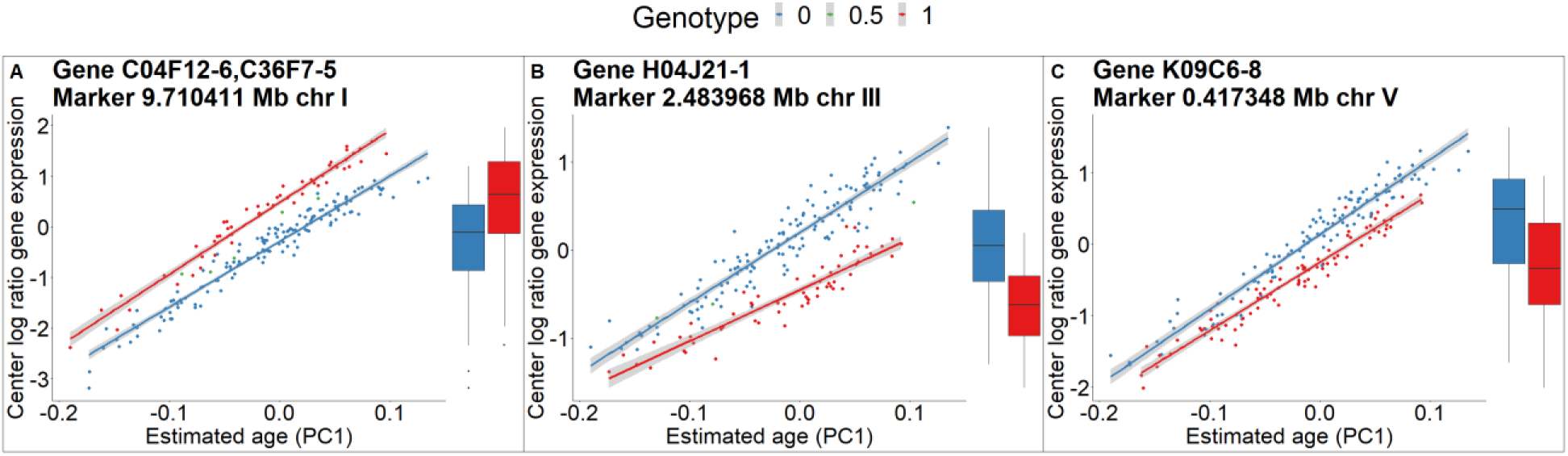
Gene expression of three example strong developmental effect eQTLs. Line plots show center log ratio of fpkm values (y-axis) over the developmental age (x-axis) for three examples of strong developmental effect eQTLs. Colors correspond to the genotype at the eQTL position. Boxplots show the magnitude of gene expression without the context of the developmental age.

Next, we investigated why some eQTLs are differentially detected between the SMM and AAM by comparing the markers with the lowest p-value according to each model. For the dynamical eQTLs none of the markers are significantly associated with gene expression according to the SMM. Plotting gene expression over the developmental age for the marker with the lowest p-value shows that at any developmental age (**Figure 4A, line plot**), there is no substantial difference between the two genotypes. However, the best AAM reveals a significant association, over the entire range of developmental ages of the mpRILs, between another marker and gene expression (**Figure 4B, line plot**). The distribution of the magnitude of gene expression between the alleles of these dynamical eQTLs can seem very similar when developmental age is not considered (**Figure 4B, boxplot**). For the single marker eQTLs a significant difference in the magnitude of gene expression (**Figure 4C, boxplot**) is mainly due to developmental variation between the alleles (**Figure 4C, line plot**). One genotype has a wider range of developmental ages on one end of the distribution compared to the other genotype, causing gene expression differences. For this subset of eQTLs, accounting for developmental variation in the model shows that not this marker, nor any of the others, is significantly associated with gene expression at any developmental age (**Figure 4D, right column**). In both cases of differentially detected eQTLs the SMM is being confounded by an association between the marker and the developmental age. These results show how dynamical eQTL mapping can reveal hidden associations and prevent developmental variation from being misattributed to a genetic marker or polymorphism.

**Figure 4:**
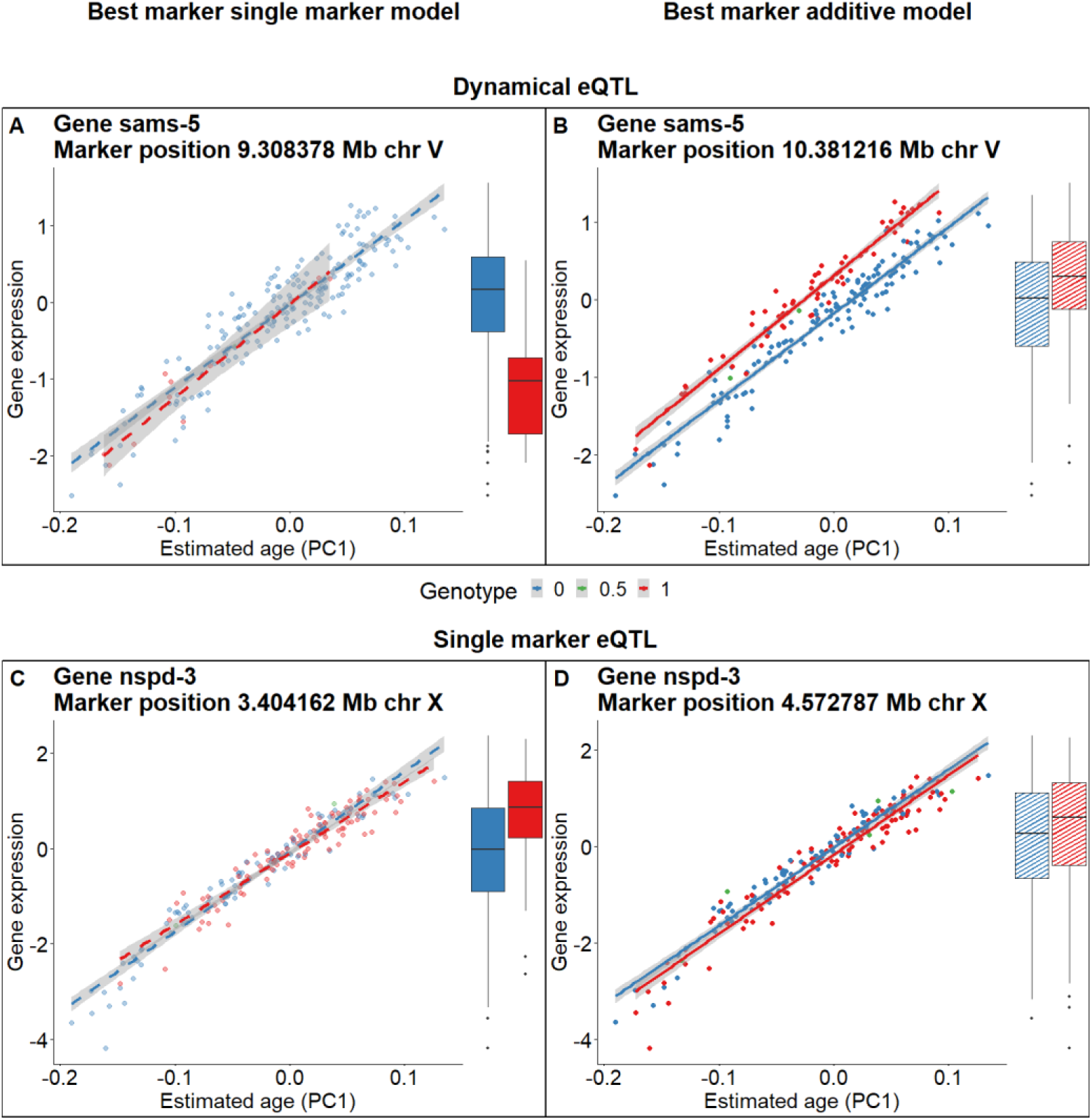
Best marker according to the SMM (A and C) or AAM (B and D) for a dynamical eQTL (A and B) and a single marker eQTL (C and D). The SMM considers only the magnitude of gene expression (solid boxplots), whereas the AAM also considers the developmental context of gene expression (solid-line plots). Dashed line-plots show gene expression over the developmental age for the best marker according to the SMM. Dashed box plots show gene expression for the best marker according to the AAM without developmental context. Line-plots are linear fits to the gene expression data. **A)** Best marker on chromosome 5 for the *sams-5* gene according to the SMM (not significant). This marker has an allele imbalance but nevertheless demonstrates the point well. **B)** Best marker on chromosome 5 for the *sams*-5 gene according to the AAM (significant). **C)** Best marker on chromosome X for the *nspd-3* gene according to the SMM (significant). **D)** Best marker on chromosome X for the *nspd-3* gene according to the AAM (not significant).

### Developmental age variation affects explanatory power of eQTLs and the narrow sense heritability of gene expression

We wanted to investigate the effect of dynamical eQTL mapping on the variance in gene expression associated with the eQTL. For some transcripts, the marker effect on the magnitude of gene expression appears much stronger after accounting for developmental variation (**Figures S7**). To quantify this, we calculated the partial eta squared of the marker variable for both the SMM and AAM. For strong developmental effect eQTLs the distribution of gene expression without considering developmental age appears much more variable within each allele (**Figure 3, box plots**). We therefore hypothesized that for these eQTLs the marker with the lowest p-value according to the SMM would have a much lower partial eta squared compared to the marker with the lowest p-value according to the AAM. Indeed, for the strong developmental effect eQTLs the distribution of the partial eta squared of the marker effect is clearly skewed towards higher values when calculated using the AAM (**Figure 5A**). We expected that the partial eta squared of the marker according to the SMM compared to the AAM could be indicative of the distribution in **Figure 2B.** Indeed, the SMM tends to infer higher partial eta squared for single marker eQTLs (**Figure 5B**), whereas the AAM tends to infer higher partial eta squared for dynamical eQTLs (**Figure 5C**). These results show that the effects of eQTLs on gene expression can be either obscured or exaggerated by hidden developmental variation, resulting in false negative and false positive eQTL detection respectively.

**Figure 5:**
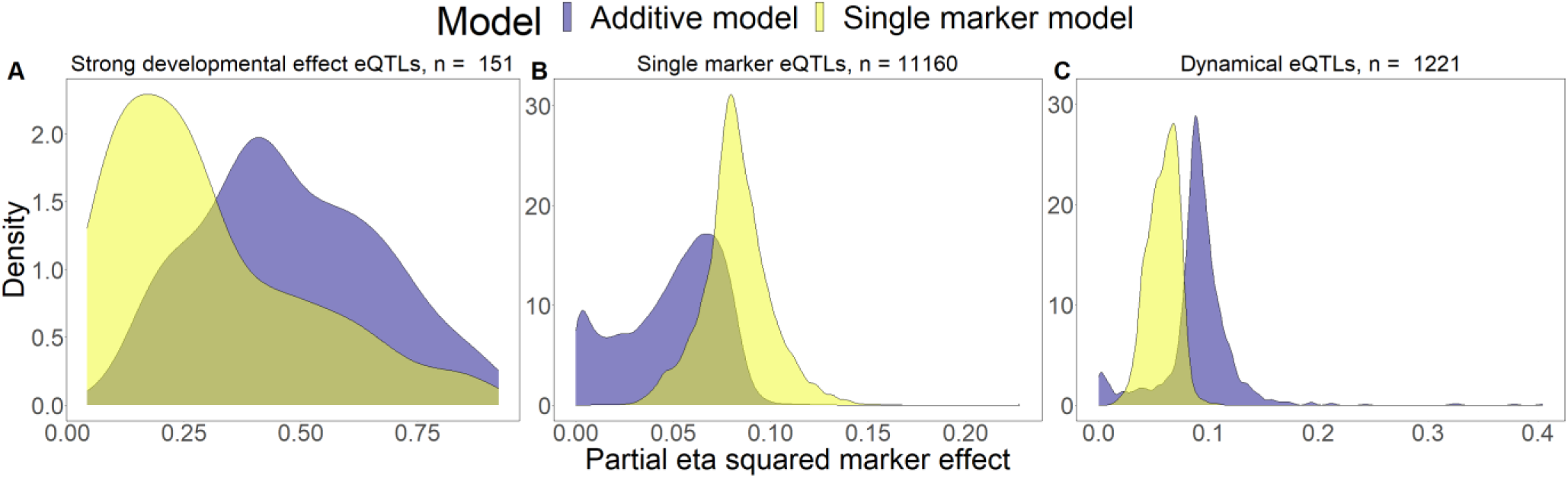
Density plot of the partial eta squared of marker effects for subsets of eQTLs according to the SMM and AAM. For the eQTLs in the subset we selected the marker with the lowest p-value according to each of the models and calculated the partial eta squared of the marker effect using the respective model. **A)** Strong developmental effect eQTLs. **B)** Single marker eQTLs. **C)** Dynamical eQTLs.

We also wanted to investigate the effect of developmental variation on the narrow sense heritability (h_2_) of gene expression. To this end, we performed a heritability analysis on the expression level of all transcripts, both with and without PC1 as a covariate. For 1210 transcripts the h_2_ increased by more than 0.05 when PC1 was included as a covariate. Conversely, for 4410 transcripts the h_2_ decreased by more than 0.05 when PC1 was included (**Table S3**). We investigated if there was an association between the type of eQTLs affecting expression and the difference in h_2_ with and without PC1 (**Figures S8**). Transcripts with no eQTLs tend to have low h_2_ with or without PC1 as a covariate, while transcripts with only eQTLs detected by both models tend to have similar h_2_, regardless of correcting for PC1. Transcripts with a larger than 0.2 difference in h_2_ when including PC1 as a covariate almost exclusively (1164 out of 1180) have either a dynamical eQTL, a single marker eQTL or both. This demonstrates that, for transcripts with eQTLs that are differentially detected between the SMM and AAM, the h_2_ of transcript levels strongly depends on variation in developmental age. The direction of the change in h_2_ also differs depending on the type of eQTL. Out of 498 transcripts with a dynamical eQTL but not a single marker eQTL, 234 transcripts have an increase > 0.05 in h_2_ when including PC1, compared to 34 transcripts with a decrease > 0.05 (**Table S3**). For transcripts with a single marker eQTL but not a dynamical eQTL the direction is reversed, with 266 transcripts with an increase > 0.05 in h_2_ with PC1 as covariate and 3092 transcripts with a decrease > 0.05.

Since eQTLs can be indicators of genomic and therefore heritable regulation of gene expression, the h_2_ of gene expression is expected to increase with the number of eQTLs. This expectation clearly holds in our data if the h_2_ is not corrected for the developmental age (**Figure 6A**). However, including PC1 as a cofactor in the heritability calculation results in lower heritability for many transcripts (**Figures S8**). Such transcripts tend to have many single marker eQTLs (**Figure S8, compare Figure S9A and S9B**), which result from developmental variation in the population rather than a direct effect of the genomic background (**Figure 4C**). Therefore, we hypothesized that the positive relationship between the number of eQTLs and the h_2_ should be reduced if the h_2_ is corrected for the developmental age. This is indeed what we observe for the SMM (**Figure 6B**). However, because eQTLs detected by the AAM do not depend on developmental variation (**Figure 4B**), the positive relationship between the number of eQTLs by the AAM and the h_2_ of gene expression is robust to developmental correction of the h_2_ (**Figure 6C, D**).

**Figure 6:**
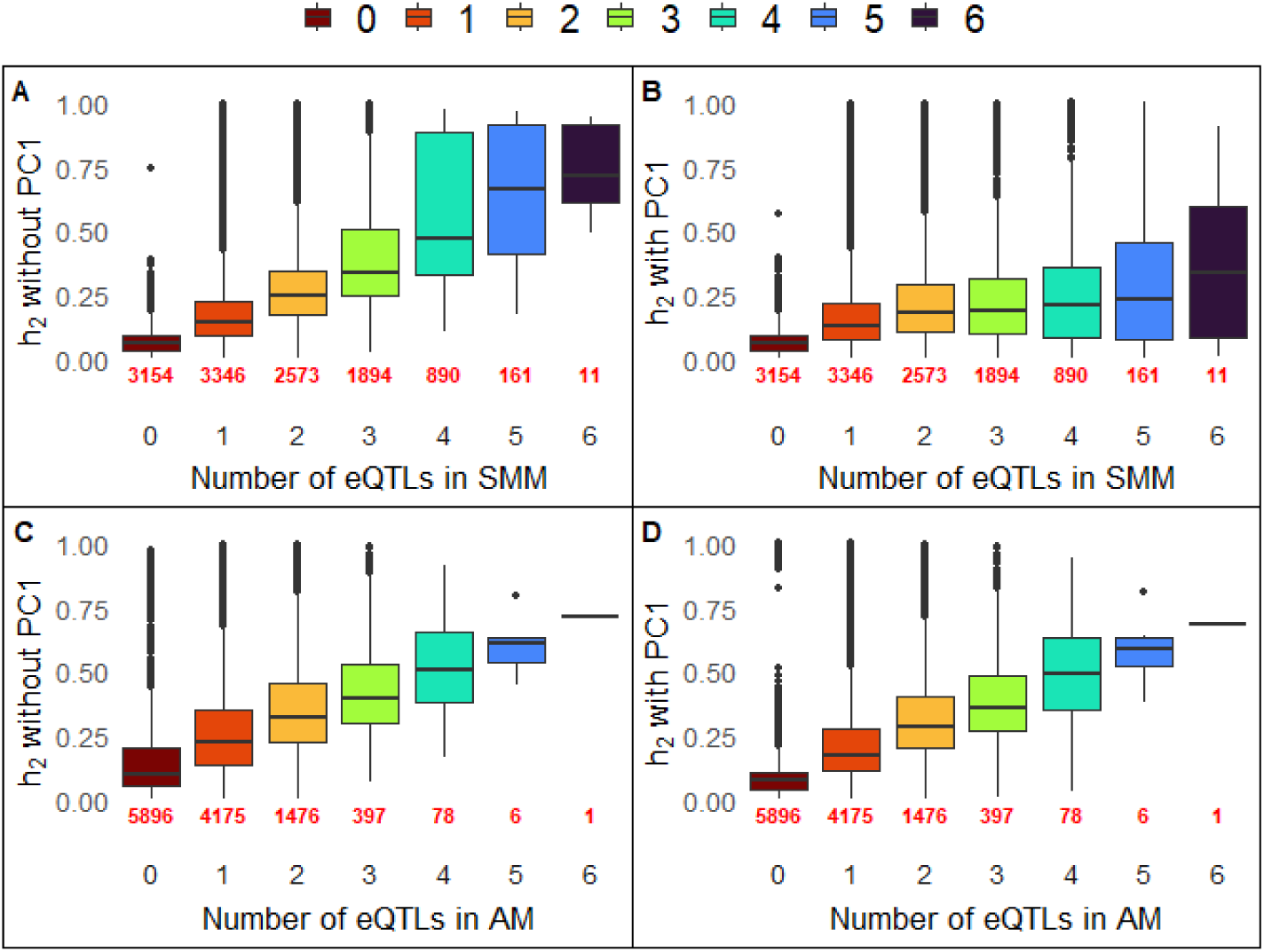
R**e**lationship **between the number of eQTLs and the h_2_.** In each subfigure we show the distribution of the h_2_ for transcripts with 0 to 6 eQTLs. Between the subfigures we show how these distributions change if we correct for development in the mapping, in the heritability calculation itself or in both. **A)** Relationship between the number of eQTLs detected by the SMM and the h_2_ of gene expression. **B)** Relationship between the number of eQTLs detected by the SMM and the development corrected h_2_. **C)** Relationship between the number of eQTLs detected by the AAM and the h_2_. **D)** Relationship between the number of eQTLs detected by the AAM and the development corrected h_2_.

### Developmental age variation affects the manifestation of eQTL hotspots

Next, we investigated whether dynamical models show an altered distribution of eQTLs over the genome compared to the SMM. The *cis/trans* distribution of eQTLs obtained with the SMM was as expected, showing several hotspots enriched in *trans-*eQTLs, whereas *cis-*eQTLs are distributed more evenly over the genome (**Figures S10**). Some of these hotspots affect the expression of hundreds of transcripts (**Figure 7A**). The largest hotspot is located at position 3.4 Mb on chromosome X and significantly affects the expression of 1196 transcripts according to the SMM. After inclusion of the developmental age in the model just a single eQTL remains at this location (**Figure 7B**). In accordance with the higher *cis/trans* ratio (**Table 1**) and the higher fraction of unique markers (**Table S2**) for eQTLs detected by the AAM, most of the other large hotspots also disappeared when we applied dynamical models.

**Figure 7:**
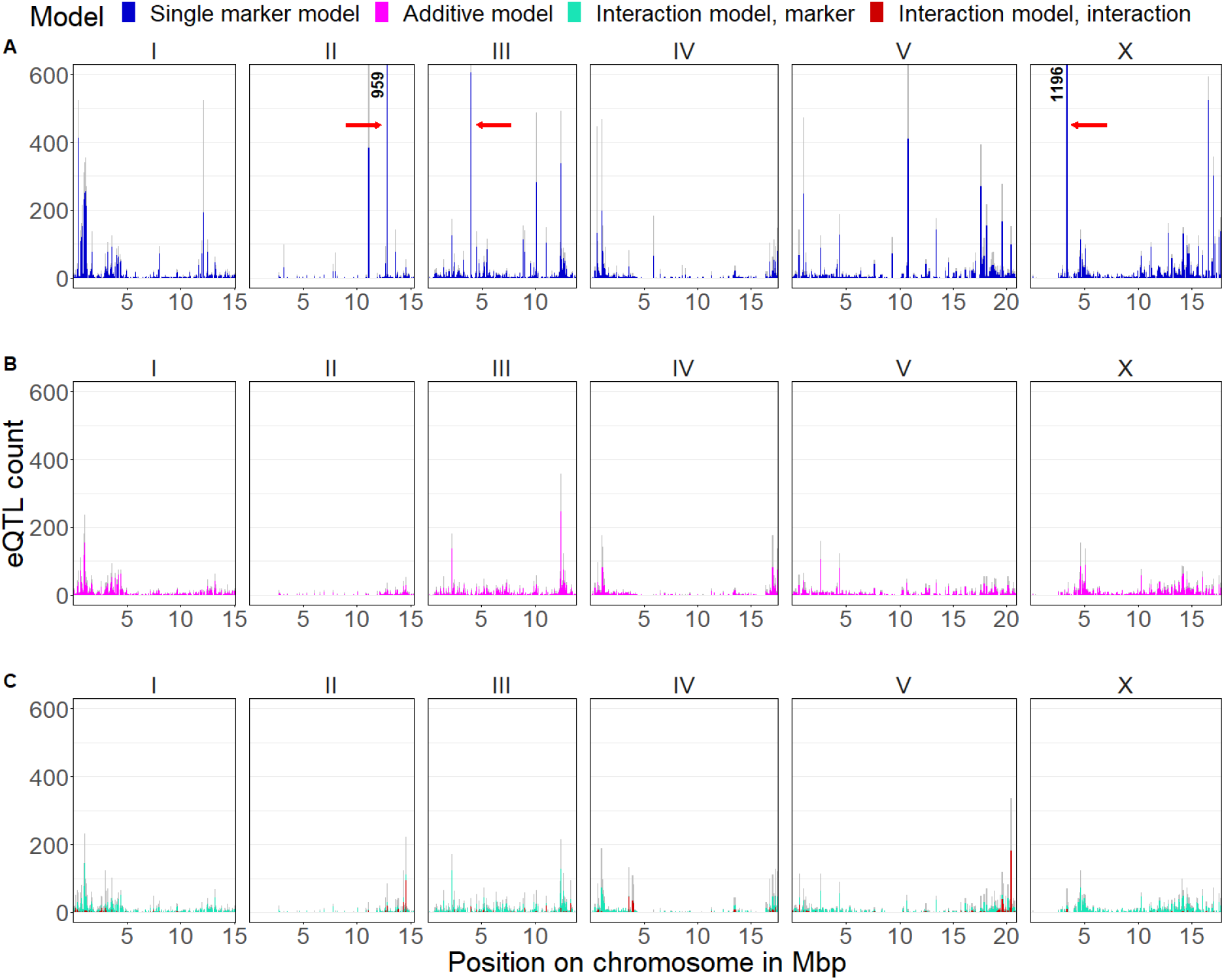
Distribution of eQTLs over the chromosome according to the different models. X-axis shows the position on the genome. Y-axis shows the number of eQTLs that map to this position. **A)** SMM. Red arrows indicate the three largest hotspots. **B)** AAM. **C)** Interaction model. Cyan and red show eQTLs according to the IMM and IMI respectively. Counts are in terms of number of transcripts mapping to the location.

We hypothesized that the difference in the distribution of eQTLs could result from a shared association between markers and gene expression with the developmental age. Under this hypothesis we would expect (1) that genes involved in developmental processes are enriched at the hotspots that disappear when applying a dynamical model, (2) that the genotype at the hotspot position is linked to the developmental age and (3) that single marker eQTLs primarily map to such hotspots.

We first investigated the enrichment of genes with developmental functions by taking the largest hotspot as an example and performing GO-term enrichment on the genes mapping to this position. The genes having an eQTL at the hotspot position were significantly enriched for GO terms related to developmental processes such as oocyte maturation, polar body extrusion after meiotic division, P granule, eggshell formation, pseudopodium, amoeboid sperm motility, male meiosis chromosome segregation, mitotic spindle pole and structural constituent of cuticle. This shows the link with development for this hotspot locus. Other hotspots were similarly enriched for genes associated with developmental processes (**See supplementary file 1**). Apart from functional categories hotspots also share many of the same genes. As an example, more than 80% (896 out of 1115) of the transcripts that map to the hotspot at position 12.8 of chromosome II also map to the hotspot at position 3.4 of chromosome X. This paints a picture of genes involved in developmental processes indiscriminately mapping to hotspots due to a shared association with the developmental age.

Second, we investigated whether the markers at the hotspots that are only found with the SMM are linked to the developmental age by comparing the distributions of the developmental age between the marker genotypes of the three largest hotspots (indicated by red arrows in **Figure 7**) (**Figure 8**). As expected, the distribution of developmental ages was clearly different between the genotypes at these hotspots. We reasoned that the markers at the positions of disappearing hotspots could be linked to the developmental age because they regulate or affect the developmental speed. In this view, such markers might affect gene expression indirectly, through their effect on the developmental process itself. To investigate we ran single marker models with the developmental age as trait. In line with our hypothesis the marker of the largest hotspot was also the most predictive of the developmental age of the mpRILs (**Figures S11**).

**Figure 8:**
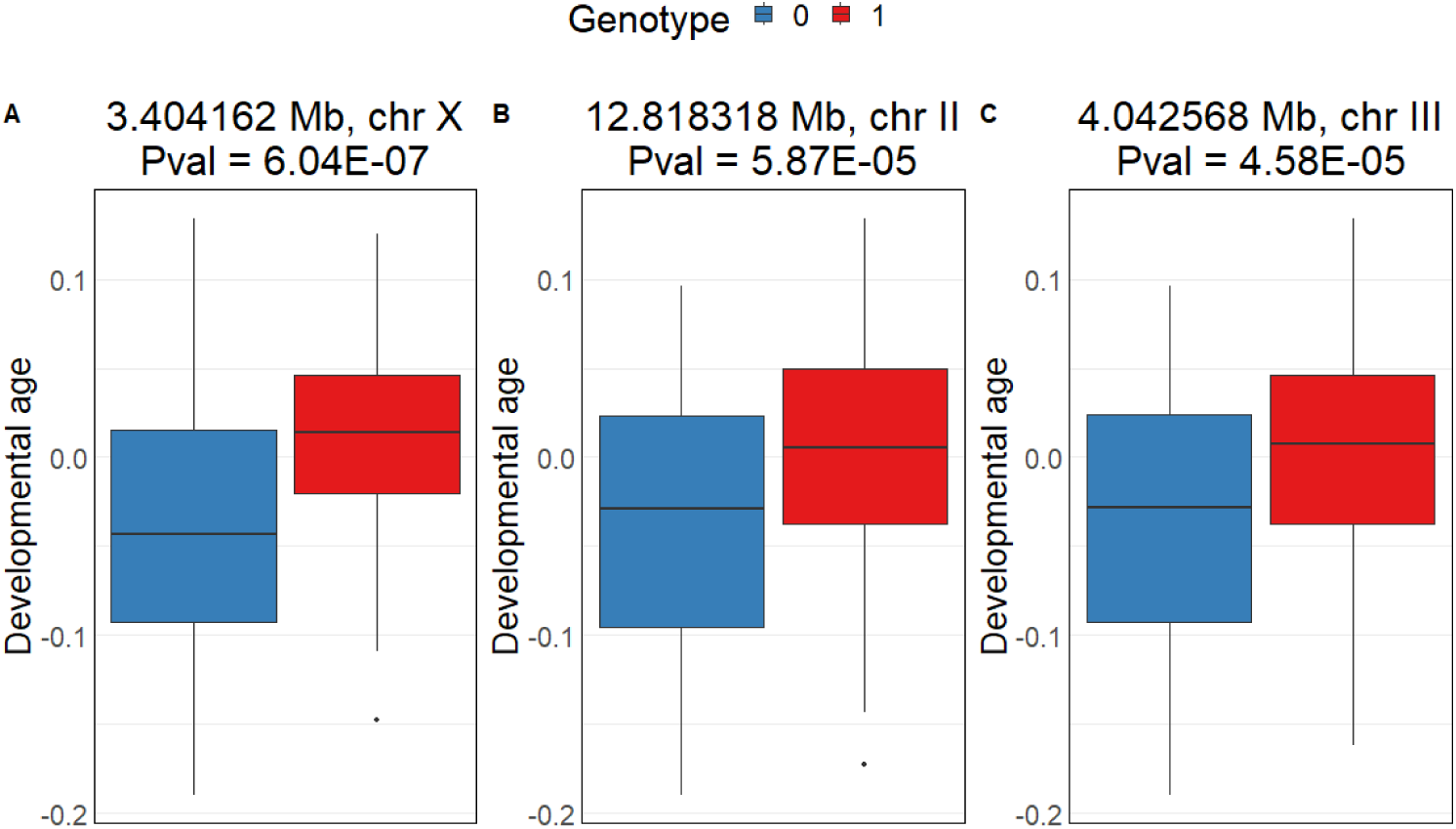
Distribution of developmental ages per allele at the three largest hotspots. Hotspots are indicated in figure 7 by red arrows. P-values are obtained with single marker linear model of PC1 ∼ marker genotype (see **Figures S11**).

Third, we investigated the distribution of single marker eQTLs (**Figures S12**). For this subset of eQTLs the best markers according to the SMM correspond to only a few loci. After adding the developmental age as an additive effect to the model, the best markers for these eQTLs are more evenly spread over the genome. For such eQTLs including the developmental age in the model controls for the shared association with the developmental age, changing significance and redistributing the markers with the lowest p-value over the genome. Together these results suggest that markers that affect the developmental speed can, as a result of the global impact of the developmental process on gene expression, be linked to the expression of many genes by non-dynamical models, which lack the developmental age as an explanatory factor (**Figure 9**).

**Figure 9:**
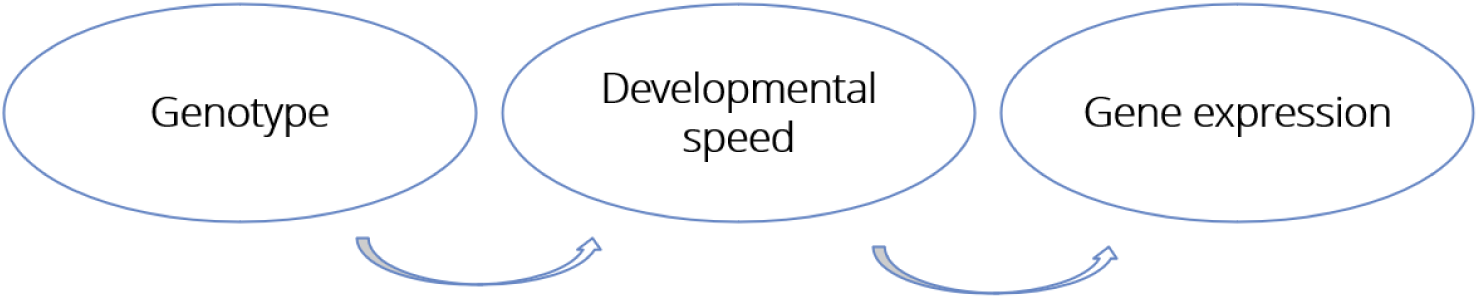
Schematic of how some hotspots cause gene expression variation. Rather than directly regulating gene expression some hotspots influence the developmental speed, causing the transcriptional program of the genotypes to be at different stages at a given moment in chronological time. This in turn causes expression variation in genes whose expression level depends on the developmental age. The developmental age is the mediator through which these loci affect gene expression.

### Interaction hotspots affect the rate at which expression is up- or down regulated during development

Genetic loci can affect not only the magnitude or timing of gene expression, but also the shape of the dynamical pattern^2^. To investigate the frequency of interactions between marker effects and the developmental age we applied a linear interaction model for eQTL mapping. A significant interaction term in this model indicates disparate dynamics between the marker alleles (**Figure 10**). Using the interaction model results in a ∼20% decrease in the number of marker effects detected (6886) compared to the AAM. On top of the marker effects the interaction model detects a significant interaction between the marker and developmental age for 1161 transcripts (FDR = 0.05). For detecting interactions, a more lenient threshold could be considered (FDR = 0.1), as this results in many additional visually convincing interactions (**Figures S13**). Investigating the distribution of interaction eQTLs over the genome shows that these also cluster in hotspots. The hotspot with the most eQTLs (20.440891 Mb, chr V) affects the expression of almost 200 transcripts. Despite the correction for the developmental age inherent to the model the genes mapping to this hotspot are significantly enriched in GO-terms associated with the cell cycle, meiosis, mitosis and other developmental processes (see **Supplementary file 2**) This locus could therefore be an important determinant in the developmental speed during the range of developmental ages spanned by the mpRILs, by affecting the rate of up- or down regulation of many genes involved in developmental processes (**Figure S14 A-C**). The genotypes corresponding to this marker also differ substantially in their distribution over the developmental age (**Figure S14D**), suggesting that its effect on the developmental speed was also present at younger developmental ages. This shows that the interaction model can detect loci that have large effects on the transcriptome by influencing the rate at which transcript levels change for many genes.

**Figure 10:**
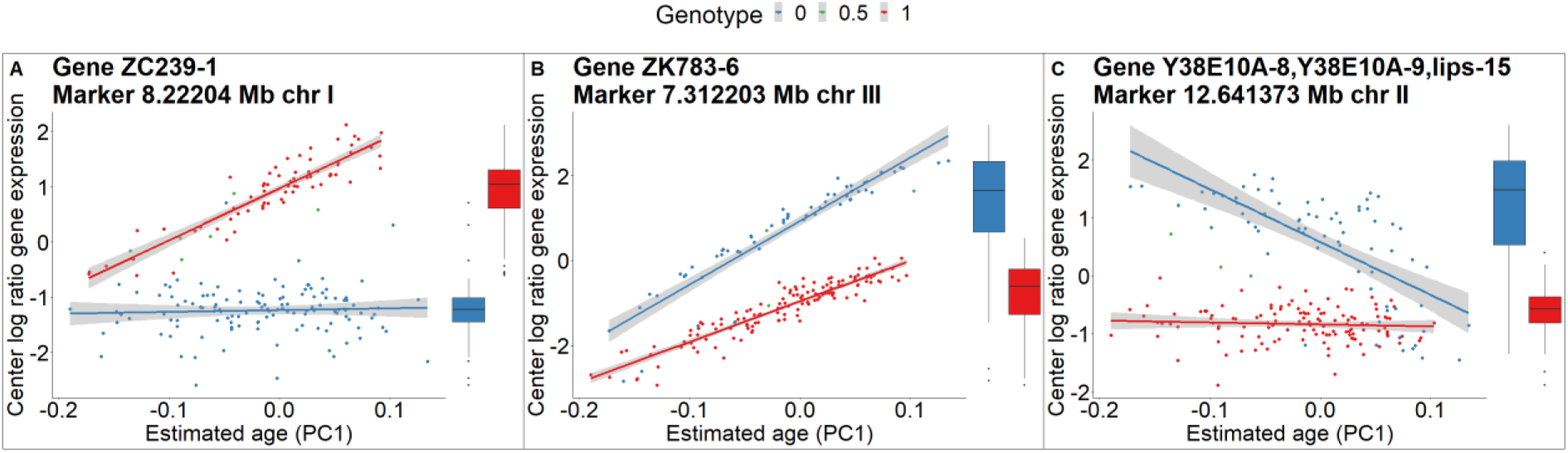
eQTLs with a significant IMI term. Colors correspond to genotypes at eQTL position. Lines are the best fit of a linear model (grey area is 95% CI) to gene expression over developmental age. Boxplots show the magnitude of gene expression without the context of the developmental age. **A)** eQTL that, depending on genotype, causes either increasing gene expression as development progresses or constant developmental dynamics. **B)** eQTL that causes the slope of gene expression over developmental age to be higher for one genotype compared to the other. **C)** eQTL for which one genotype exhibits negative regulation of gene expression while the other genotype shows constant gene expression. In this case, the linear interaction model is a good enough approximation of a gene expression pattern that is non-linear. Note that the rate at which expression levels of this transcript are decreasing for mpRILs with the blue allele increases with increasing developmental age.

## Discussion

We have shown that eQTL hotspots that disappear after controlling for the developmental age can correspond to markers influencing the developmental speed. For such hotspots, there is likely no direct regulatory relationship between the hotspot and the genes with an eQTL mapping to it. After all, at any specific developmental age there is no significant difference in expression between the marker genotypes. Rather, one of the genotypes is on average further developed than the other, causing gene expression differences. Because the genes mapping to such hotspots are enriched for developmental functions, they are likely to have non-constant expression over the developmental age range of the mapping population. A non-zero slope of gene expression over the developmental age can cause differences in the overall magnitude of gene expression between marker alleles, even in the absence of gene expression differences between marker alleles at any specific developmental age, if developmental ages are not distributed equally between the alleles.

The mechanism through which a polymorphic SNP influences the developmental speed can be difficult to entangle because gene expression both drives and is the result of the developmental speed. Therefore, an eQTL might influence the developmental speed by affecting the expression of some genes, as a result changing the expression of many more genes. For some SNPs in hotspots the association with the developmental age can be understood from the genomic context. For example, the large hotspot at the beginning of chromosome X maps close to the *vit-5* gene encoding for the Vitogellin-5 protein which is a major yolk component. Inhibition of *vit-5* by means of RNAi has been shown to result in slower rates of post-embryonic growth^30^. Interestingly, the maternal expression of vitellogenin genes appears to be a major determinant of the developmental speed^26^. As a second example, the hotspot at 10.1 MB on chromosome III is located in the *kel-10* gene. The human ortholog (*KLHL10*) of this gene is involved in spermatogenesis, and *kel-10* is affected by the *daf-2* gene, a well-known regulator of lifespan in *C. elegans*^31^.

Hotspots are often assumed to be global regulators of gene expression, although it is well known that hotspots can also result from genes with correlated expression because of uncontrolled latent factors^32, 33^. We show that developmental variation can be a major cause of such correlations, to the extent that most prominent hotspots are no longer present after correcting for the developmental age. Therefore, we posit that the default interpretation for hotspots should be an association between the hotspot locus and the developmental age.

Loci that influence the developmental speed need not be seen as merely a confounder in eQTL mapping. Instead, they are crucial determinants of the expression state of an organism. Investigating the eQTLs detected by a non-dynamical but not a dynamical model is a convenient approach to search for loci that are causative for developmental variation. This would allow a distinction between loci that cause gene expression differences even if organisms are at the same developmental age (eQTLs detected by AAM), and loci that affect gene expression by causing developmental age differences (single marker eQTLs). This approach is not limited to the developmental process but can be applied to any process which leaves a sufficiently strong signal in transcriptomics data. More generally, including latent factors in eQTL mapping can allow one to distinguish between loci that influence gene expression through their effect on generic processes and direct regulators of gene expression.

We calculated the narrow sense heritability (h_2_) of gene expression both with and without correction for the developmental age. Comparing the standard h_2_ with the development corrected h_2_ could provide an indication of whether a gene’s expression is in part heritable through the genetical component of developmental age differences, or mostly independent from developmental age. In the first case one would expect the development-corrected h_2_ to decrease compared to the standard h_2_. On the other hand, if the heritability of gene expression is independent from the developmental process, one would expect the development-corrected heritability to be similar or higher compared to the standard h_2_. While the developmental age is itself heritable and can thus have a real effect on the heritability of expression, it can also confound heritability analysis. This is evidenced by the subset of transcripts with a much larger development corrected h_2_ compared to standard h_2_ (**Figures S8, S9**). Possibly, this confounding effect can be explained by the relatively low h_2_ of PC1 (∼0.23), our proxy for the developmental age. A large part of the variation in developmental age could be due to stochastic factors such as small fluctuations in temperature within the experimental setup^34^ or non-additive genetic effects. This part of developmental variation could obscure the effect of the genomic background on the narrow sense heritability of gene expression for some transcripts, resulting in more accurate heritability estimates when controlling for development.

In this paper we used a relatively simple method (PCA) to quantify the developmental age. Various more complex methods that extract developmental ages from transcriptomics data have already been developed. An example is the RAPToR framework, which projects the samples on an interpolated reference time series^23^. Because we wanted a method that was generally applicable, we did not want to be completely dependent on a reference (time) series. Our approach enables detection of other (semi-)linear processes affecting gene expression on a genome-wide scale and can be applied to other published eQTL studies in *C. elegans*^35–40^ and beyond^27^. Furthermore, our method, despite being simpler, provided an ordering of the expression of developmental indicator genes that was more consistent with a monotonic linear increase.

Because eQTLs can also cause allele dependent non-linear expression patterns over the developmental age^2^, we performed the eQTL mapping using a natural spline of the developmental age (**Supplementary text A1**). Using the interaction term of this model, we hoped to identify eQTLs responsible for complex dynamical patterns. However, constraining the FDR of the natural spline model using the same permutation strategy we used for the linear models was not possible, as the p-values obtained on the actual dataset were barely distinguishable from the p-values obtained on the randomized dataset. We suspect this is in part due to a lack of statistical power, resulting from a combination of the size of the population (199 lines) and the dense genetic map (8933 SNPs) used for the mapping. An additional factor could be that our population has a narrow distribution of developmental ages. A study by Francesconi and Lehner detected many non-linear patterns using natural splines^2^. The population used in their study contained worms in the L3, L4 and young adult developmental stages. The mpRILs used in this study were all in the L4 larval stage (**Figures S15**). Possibly, non-linear dynamics only occur over a wider range of developmental ages than is covered by the mpRILs in our study or when multiple distinct stages are present. To investigate we clustered 2346 transcripts whose expression oscillates with a period of approximately eight hours^20^ using k-means. The clustering revealed mostly monotonic or constant gene expression patterns for the clusters (**Figures S16**). This indicates that the developmental range spanned by the mpRILs is not broad enough to clearly observe the oscillations to which a large part of the *C. elegans* transcriptome is subject^20–22^. Therefore, the natural spline model may not be suitable, or at least superfluous, for mapping eQTLs in our experiment.

We showed that PC1 accounted for most of the developmental variation in our population. The study by Francesconi and Lehner showed that the developmental signature was distributed between PC1 and PC2, which were assigned the interpretation of oogenesis and spermatogenesis respectively^2^. Together PC1 and PC2 explained ∼50% of the gene expression variation, which is close to our PC1 (∼48%). In accordance with two developmental axes, the projected samples formed an almost circular trajectory on PC1 and PC2. The authors showed that, while the distribution of developmental ages is centered on the L4 larval stage, the population also includes individuals from late L3 and early adult. It is possible that transitions between developmental stages cause developmental variation in gene expression to be distributed over more than one PC axis.

A previous study scored the time to first egg phenotype of the mpRILs^24^. Because time to first egg is a clear indicator of the developmental speed, we hypothesized that this phenotype would correlate with PC1. Surprisingly, there was almost no correlation between PC1 and the time to first egg, or indeed any of the other phenotypes scored in this study (**Figures S17**). Considering the relatively low h_2_ of PC1 (∼0.23), this can likely be partially attributed to between experiment variation. A second relevant factor could be the observed inverse relationship between the duration of larval development and the time between the first adult molt and the development of the first embryo^26^. Thirdly, the developmental speed could differ between developmental stages. Development is a complex process with crucial developmental events likely being regulated by independent timers^41^. Accordingly, correlations between the durations of two larval stages in experiments with genetically identical worms are low when temperature is strictly controlled^34^. On top of this, the genetic background can have stage-specific effects on the developmental speed^34, 41^. Stage-specific developmental variation would de-correlate developmental ages over the course of chronological time. Verifying this third explanation would require an experiment in which samples are obtained within and between time points. Such an experiment would allow us to measure the degree to which developmental variation is maintained on various timescales. More importantly, this experiment would allow us to compare gene expression over developmental age with gene expression over chronological age. In this way we could definitively show whether gene expression variation over PC1 is due to developmental variation. Furthermore, it would allow for an investigation into the extent to which eQTL patterns over the developmental age translate to eQTL patterns over chronological time.

In conclusion, we have shown that performing a PCA and including principal components into the model for eQTL mapping leads to an improved understanding of eQTL patterns by separating differences in gene expression due to variation in generic processes from direct effects of the genetic background. Furthermore, the effect of these generic processes on gene expression and their interaction with the genetic background can be understood by comparing models with and without principal components. Within an experimental condition, principal components can correspond to the developmental process, technical noise or other latent variables unrelated to the research question. Including principal components as representations of such latent variables in the mapping procedure is a convenient way to control for or come to understand sources of variance confounding the effect of interest^42^. We recommend that this should be standard practice in eQTL mapping studies given the large effect of processes like development on the transcriptional state of the cell.

## Methods

All analyses were performed using R version 4.1.3^43^. Plots were generated using the ggplot2 package^44^. Pearson correlations were calculated using the cor() function from base R.

### Data and pre-processing

We used data of 199 multi-parental recombinant inbred lines (mpRILs)^24, 25, 29^. The mpRILs were derived from an advanced cross between four parental lines which were collected from the French regions of Orsay (2 lines) and Santeuil (2 lines). The mpRILs were grown for 48 hours after bleaching at 24 degrees. We verified that our population was still in L4 by calculating the correlation of the transcriptome of the mpRILs with two L4 reference genomes and two young adult reference genomes from the N2 strain (L4, L4b, YA and N2Yad-1 from ^6^) (**Figures S15**). The parental lines were excluded from the eQTL mapping procedure and all subsequent analysis unless explicitly mentioned. The data consists of fpkm values for 38322 transcripts obtained using RNA-seq and a genetic map of 8933 bi- allelic SNPs. Due to the SNPs being bi-allelic the genotype at each marker was coded as 0 or 1 for homozygous individuals and 0.5 for heterozygous individuals. Before analysis we filtered the gene expression data to retain the 12029 transcripts with more than 20 non-zero fpkm values and a log2(mean fpkm) value higher than -5. Out of the 12029 filtered transcripts 2341 are polycistronic, for a total of 15224 genes in the filtered dataset. In all analysis we use the center log ratio of gene expression (CLR), which for gene j and mpRIL i is calculated according to the following equation:

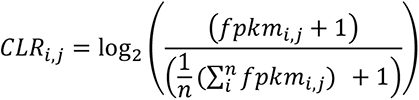

### Developmental age

We quantified the developmental age by performing PCA on the filtered and normalized gene expression matrix (with the mpRILs as columns and the transcripts as rows, such that mpRILs have loadings on PC1 and transcripts have scores) using the prcomp() function from the stats package. The loadings of the mpRILs on PC1 were taken as a proxy for the developmental age in subsequent analysis. For figure 1, S2, S3, S4 and S12 we calculated the Z-score of the center log ratio of the fpkm values by mean centering on 0 and scaling such that the expression of each gene has a standard deviation of 1 using the scale() function from base R.

We calculated developmental age estimates using the ae() function from the RAPToR package in R. We tried both the Cel_larval and Cel_larv_ya reference sets, obtaining a more even spread in distribution using the former. The ae() function allows for the possibility of specifying a prior but this did not provide a narrower distribution of the estimated ages, even when using a low standard deviation (1 hour). We therefore estimated the ages of the mpRILs with the Cel_larval reference and no prior.

### eQTL mapping with linear models

We ran linear models using the FastLm() function from the RcppAramadillo package. We applied a single marker model (gene expression ∼ marker genotype), AAM (gene expression ∼ marker genotype + PC1) and interaction model (gene expression ∼ marker genotype (IMM term) + PC1 + marker genotype * PC1 (IMI term)) to perform pairwise tests between the filtered transcripts and all markers. P-values were calculated for model terms using t-tests. For the interaction model, both a significant IMM term and a significant IMI term were considered an eQTL. Per gene, we called a maximum of one eQTL per chromosome. We called a *cis*-eQTL if the distance between the middle of the gene and the marker was < 2Mb, and a *trans-*eQTL otherwise. For all models, we also generated p-values by randomly distributing the gene expression values over the mpRILs for each gene. We used the p-values obtained by permutation to constrain the type I error and obtain significance thresholds. For thresholding the marker and interaction terms we used the lowest p-value per gene per chromosome. For the developmental age variable, we used the p-values of the models with the lowest marker p-value per transcript per chromosome. All results are obtained using an FDR of 0.05 unless explicitly mentioned in the text. P-value thresholds are in **Table S1**.

### Transcripts affected by development

Transcripts were considered significantly affected by the developmental process if they had a significant p-value for the development variable using the AAM, in all the six models (one per chromosome) that resulted in the lowest marker p-value.

### Calculating partial eta squared and heritability

We performed an anova on the model with the lowest marker p-value using the aov() function from the stats package and extracted from the effects vector the residual sum of squares (RSS) and the sum of squares explained by the marker (SSM). Note that the marker with the lowest p-value can differ between the SMM and the AAM. We then calculated the partial eta squared according to the following equation:

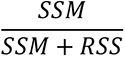

We calculated narrow sense heritability using the marker_h2() function from the heritability package. This function requires a kinship matrix as input, which we calculated using the A.mat() function from the rrBLUP package. We calculated the heritability of gene expression both with and without the loadings on PC1 as a covariate.

### GO enrichment

We performed GO-term enrichment on genes that map to specific hotspots. We tested for enrichment of GO terms from the subcategories biological process, molecular function and cellular component using a hypergeometric test. As the total gene set we used the genes on the 12029 transcripts used in the eQTL mapping. We called significant enrichment if p-value < 0.001 and at least three genes were associated with this GO-term.

## Author contributions

BLS conceived the study, BLS and BvE designed and performed the data analysis steps. BvE and BLS wrote the paper assisted by HN, JEK and MS.

## Acknowledgements

M.G.S. was supported by NWO domain Applied and Engineering Sciences VENI grant (17282).

## Supplementary tables

**Table S1:** –log10 p-value thresholds linear models.

|  | Single marker model | Additive age model |  | Interaction model |  |  |
| --- | --- | --- | --- | --- | --- | --- |
| FDR | Marker | Marker | Developmental age | Marker | Developmental age | Interaction |
| 0.05 | 4.06 | 4.44 | 1.79 | 4.60 | 3.07 | 5.60 |
| 0.1 | 3.52 | 3.87 | 1.43 | 3.98 | 2.65 | 4.52 |

**Table S2:** Statistics on quadrants, denoted by different colors in figure 2B. The first row describes the number of gene/marker combinations per quadrant, the second and third row show the number of unique markers per quadrant for the SMM and AAM respectively and the fourth row displays the number of eQTLs per quadrant for which the location of the best marker changes by at least 1MB between the SMM and AAM.

|  | Brown quadrant | Orange quadrant | Cyan quadrant | Purple quadrant |
| --- | --- | --- | --- | --- |
| Total gene/marker combination | 52348 | 11160 | 1221 | 7445 |
| Unique markers SMM | - | 1120 | 474 | 2364 |
| Unique markers AAM | - | 2300 | 676 | 2726 |
| Number of markers switching between models | - | 5306 | 667 | 1576 |

**Table S3:** Differences in heritability with and without adding PC1 as a cofactor for subsets of transcripts. Percentages are in terms of total number of transcripts in the subset (shown in left column).

| | Total transcripts | Number of genes with<br>0.05 increase in $h^2$<br>when adding PC1 as<br>covariate | Number of genes with<br>0.05 decrease in $h^2$<br>when adding PC1 as<br>covariate |
| --- | --- | --- | --- |
| All transcripts | 12029 | 1210<br>10.1% | 4410<br>36.7% |
| Transcripts with no<br>eQTL | 2842 | 347<br>12.2% | 319<br>11.2% |
| Transcripts with same<br>eQTL between SMM<br>and AAM | 2589 | 244<br>9.4% | 296<br>10.4% |
| Transcripts with<br>dynamical eQTL but<br>not a single marker<br>eQTL | 498 | 234<br>47.0% | 34<br>6.8% |
| Transcripts with single<br>marker eQTL but not a<br>dynamical eQTL | 5451 | 266<br>4.9% | 3092<br>56.7% |
| Transcripts with a<br>single marker eQTL<br>and a dynamical eQTL | 649 | 119<br>18.3% | 330<br>50.8% |

## Supplementary figures

**Figure S1:**
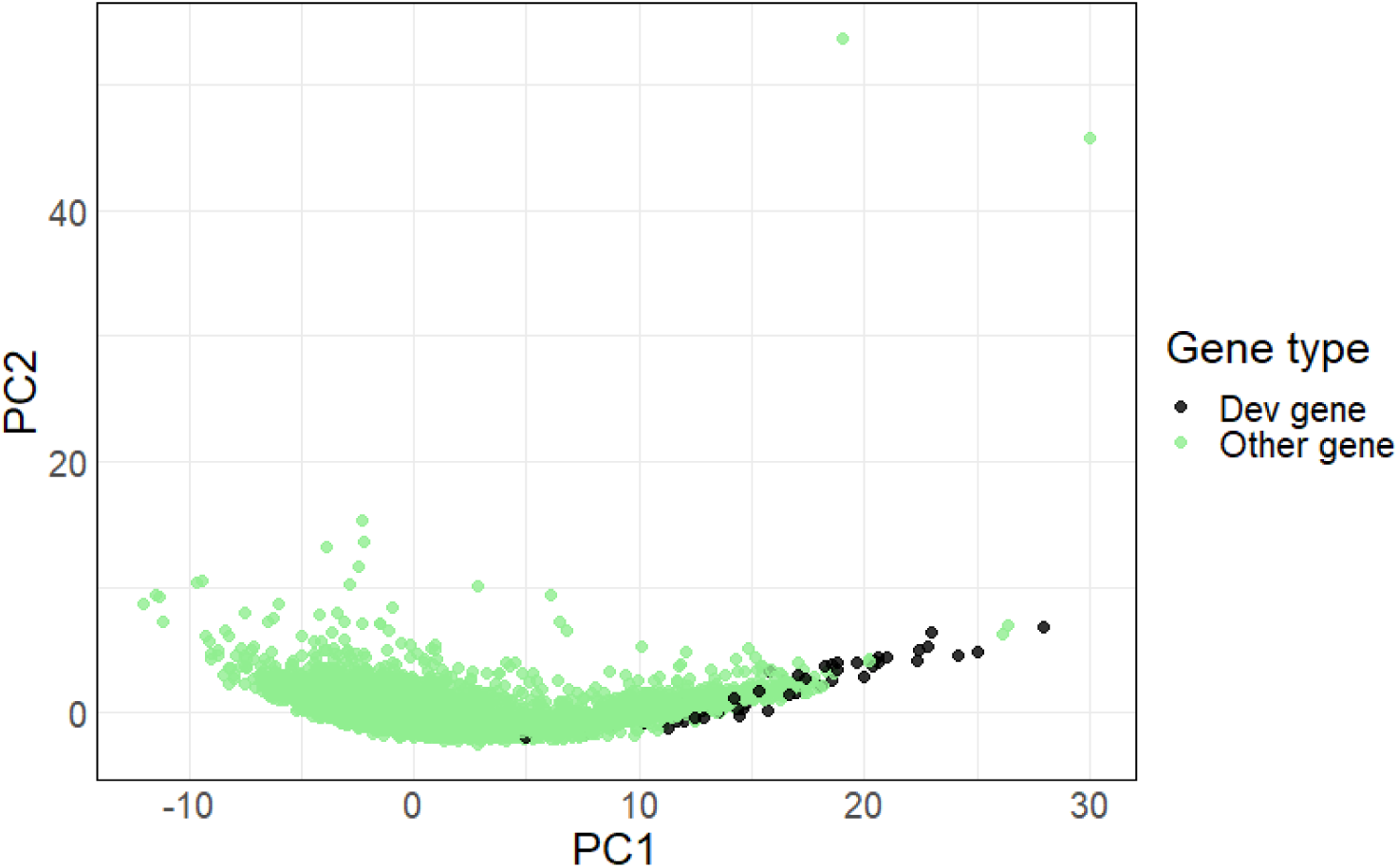
Developmental indicator genes have high scores on PC1. Projections of all 12029 genes used in the mapping on PC1 (x-axis) and PC2 (y-axis). Black color denotes the 53 developmental indicator genes, while the green color denotes all other genes.

**Figure S2:**
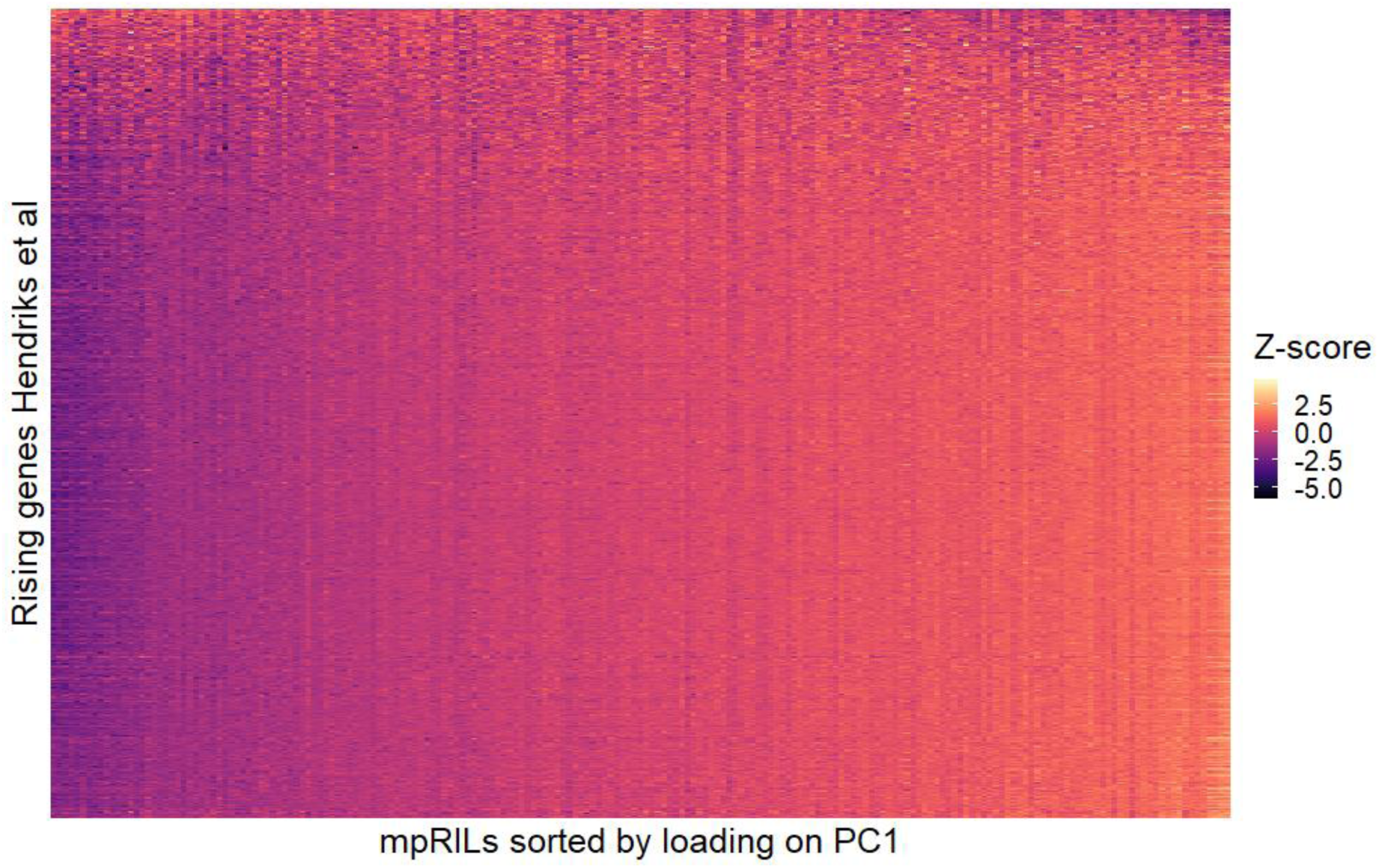
Heatmap of expression monotonically rising genes. Heatmap of z-score of the center log ratio of gene expression of 2050 genes that were identified as monotonically increasing between the beginning of the L3 stage and the young adult stage^20^. The columns are ordered by the projection of the mpRILs on PC1 and the rows are ordered from low to high by the slope of the expression values over PC1. Slopes were calculated as the coefficient of the developmental age variable in a linear model of gene expression ∼ PC1.

**Figure S3:**
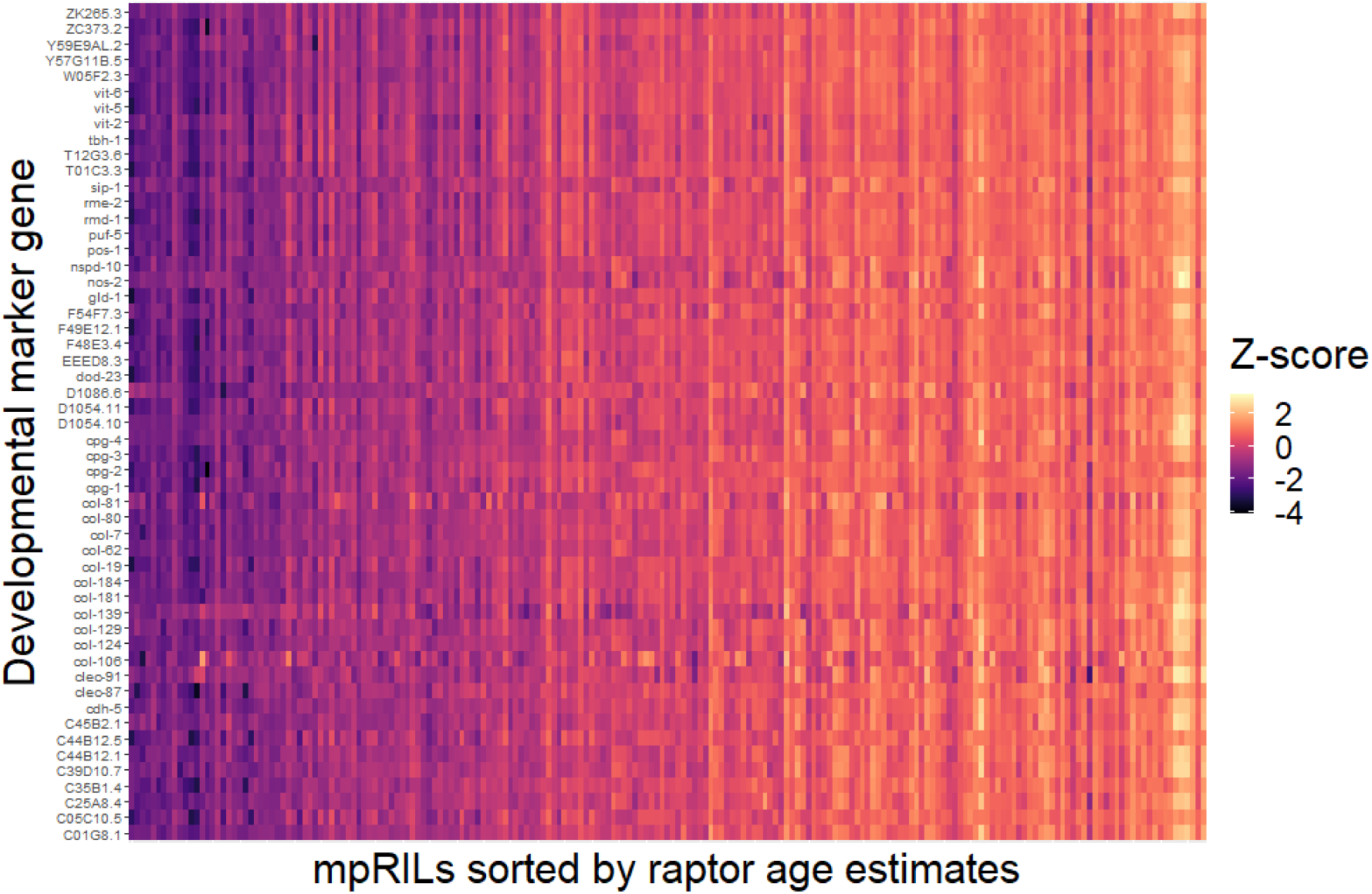
Expression of developmental indicator genes of mpRILs ordered by RAPToR age estimates. Heatmap of z-score of the center log ratio of gene expression of developmental indicator genes (y-axis) for the mpRILs sorted by the RAPToR age estimates (x-axis).

**Figure S4:**
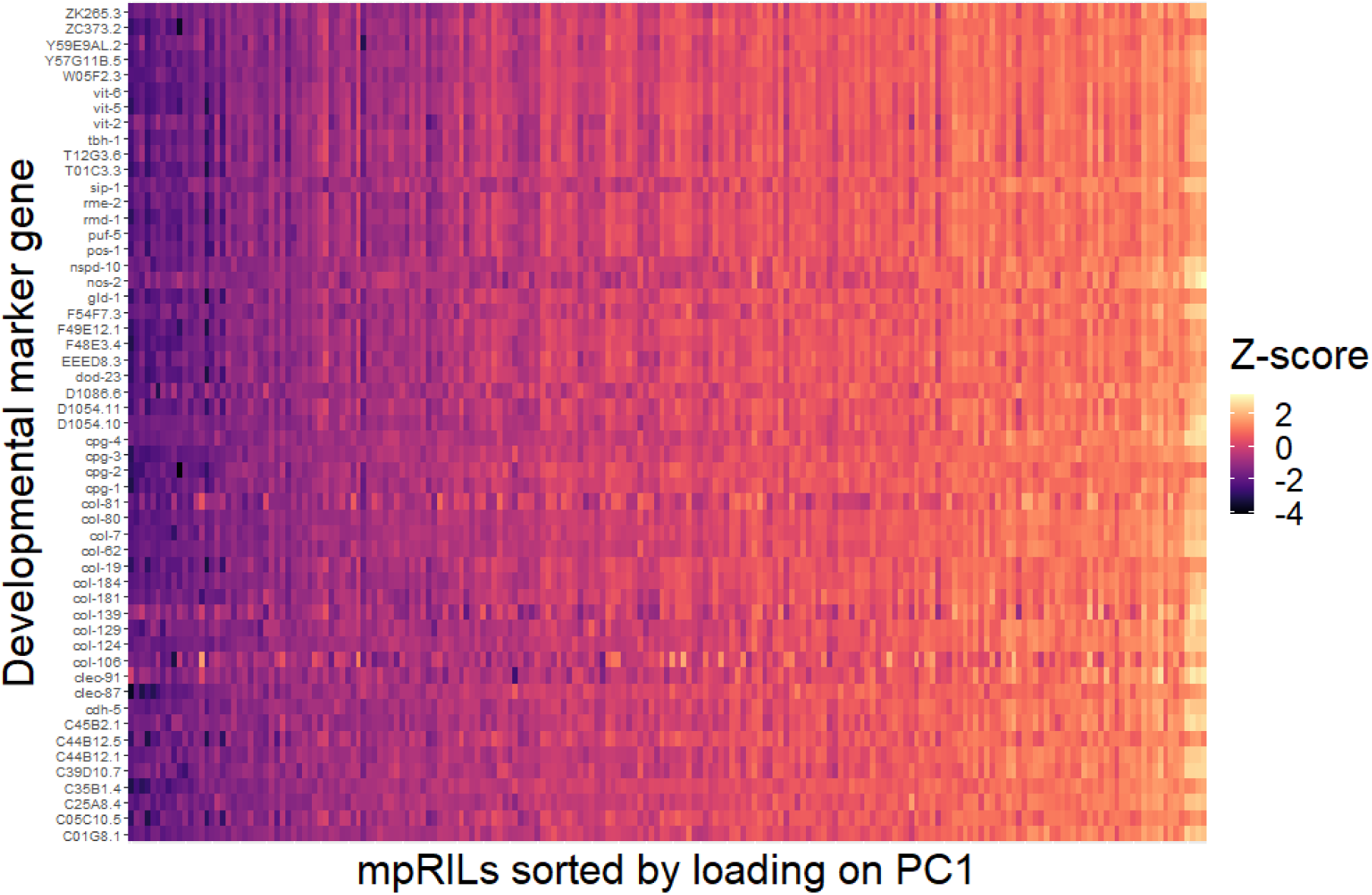
Expression of developmental indicator genes of mpRILs ordered by PC1. Z-score of center log ratio of gene expression of developmental indicator genes (y-axis) plotted for the mpRILs sorted by their projection on PC1 (x-axis).

**Figure S5:**
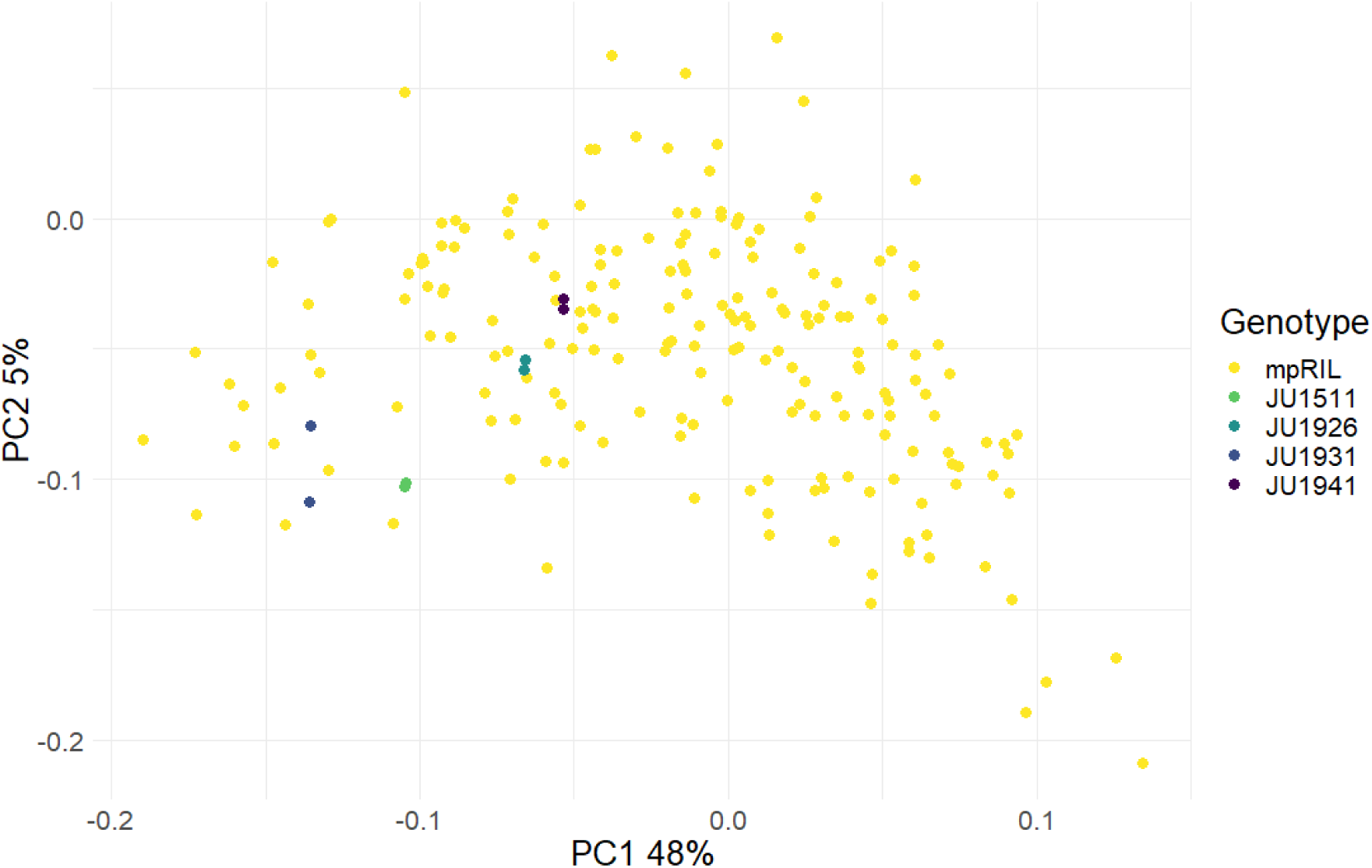
Parental replicates have almost identical projections on PC1. Projection of population on the first two principal components, colored by whether the worm corresponds to a parental genotype or one of the mpRILs.

**Figure S6:**
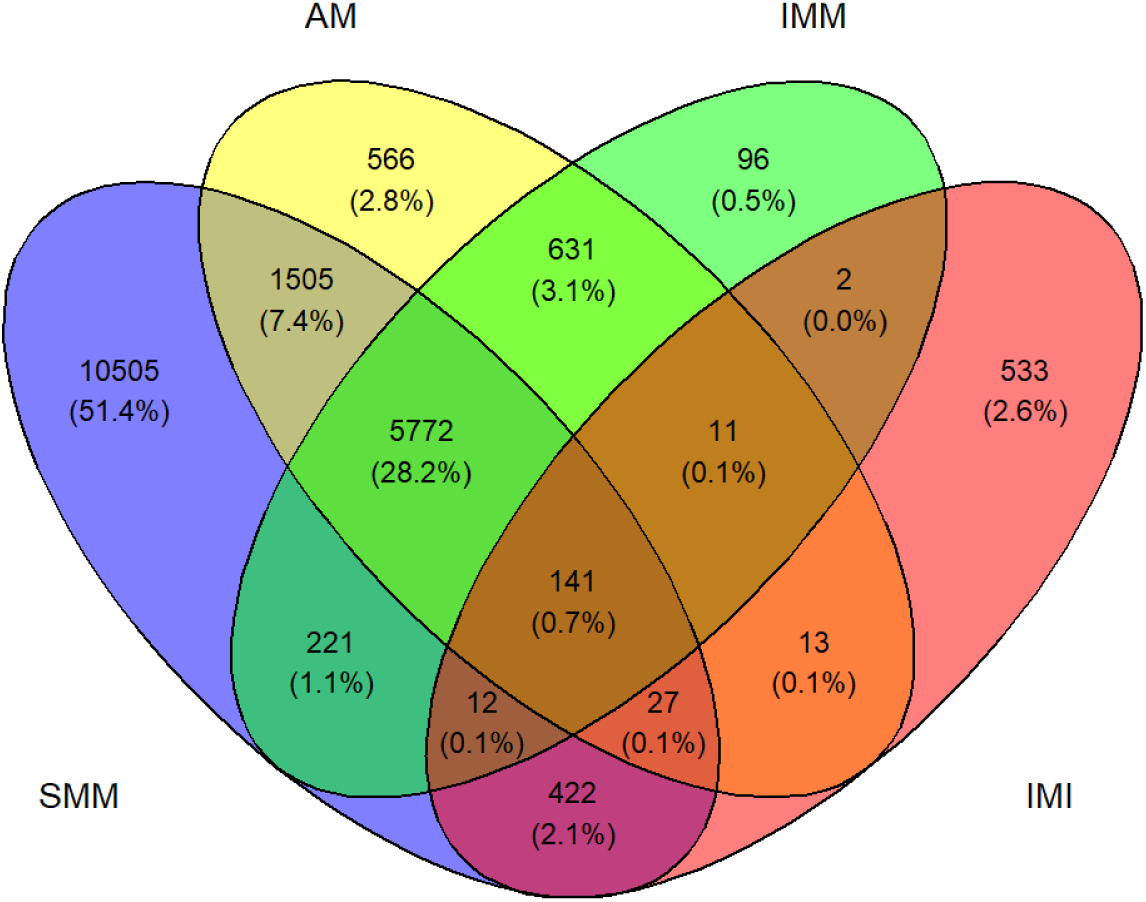
Venn diagram of eQTLs overlap between various models. Overlap between the single marker model (SMM, blue), additive age model (AAM, yellow), interaction model, marker term (IMM, green) and interaction model, interaction term (IMI, red). eQTLs are overlapping between models if the models detect a significant marker on the same chromosome for the same gene. Percentages are in terms of the union of the eQTLs detected by the four models.

**Figure S7:**
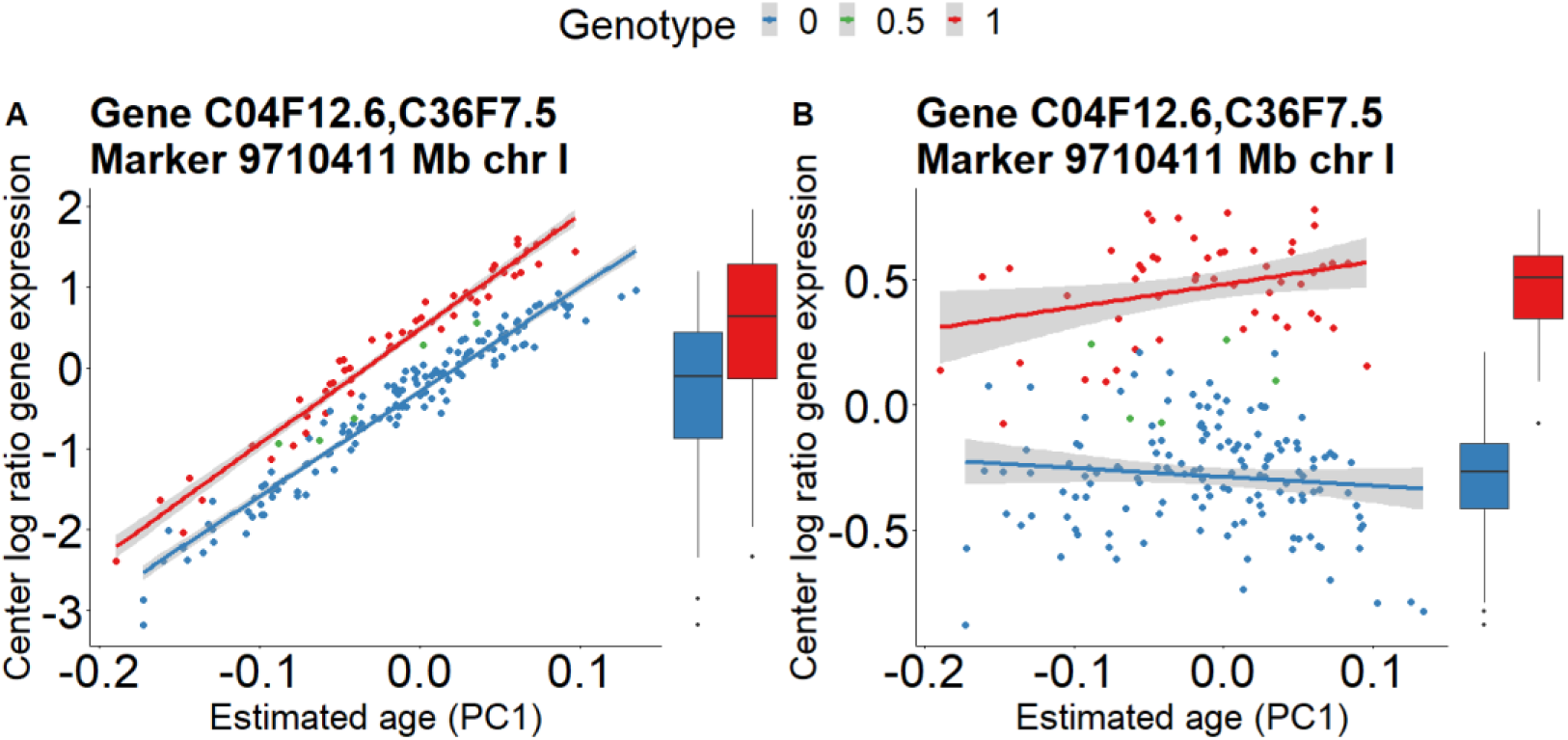
Gene expression before and after removing developmental variation for an example eQTL. Color corresponds to genotype at eQTL position. We remove developmental variation by calculating the coefficient of the developmental age variable in the additive age model. We then subtract this coefficient multiplied with the projection on PC1 from the expression of each of the mpRILs. Lines are best linear fit (grey area is 95% CI). The boxplots show that the strength of the marker effect on the magnitude of gene expression appears lower **A)** before compared to **B)** after removing developmental variation.

**Figure S8:**
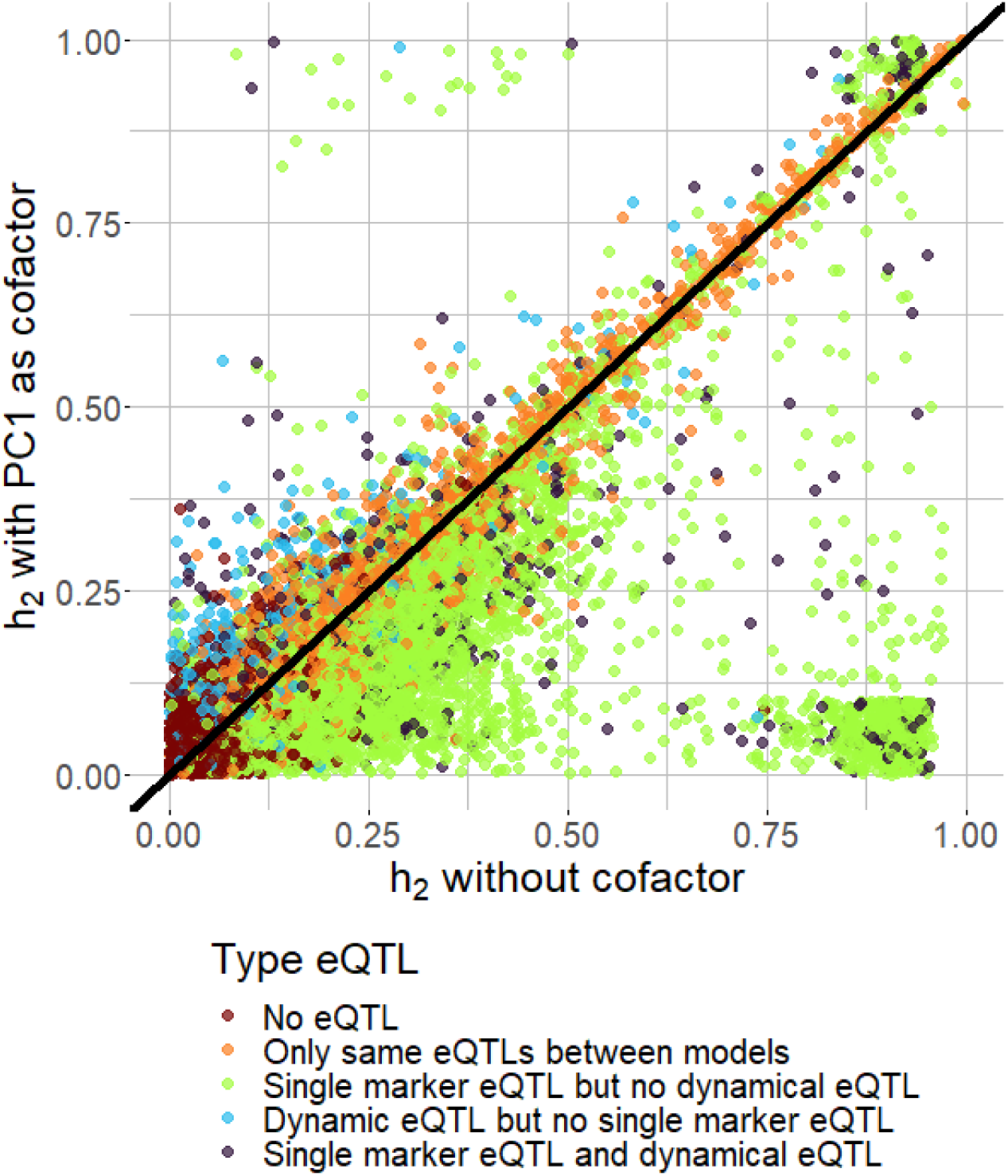
Heritability of gene expression with and without PC1 as cofactor depends on the type of eQTLs. The x-axis shows the heritability calculated without PC1 as covariate. The y-axis shows the heritability calculated with PC1 as covariate. Each dot represents one of the 12029 transcripts with expression in our dataset. Black line has slope 1 and passes through the origin. Transcripts are colored by whether they have no eQTLs according to the SMM and AAM (orange), the same eQTLs between both models (green), a single marker eQTL but not a dynamical eQTL (light blue), a dynamical eQTL but not a single marker eQTL (brown) or a single marker eQTL as well as a dynamical eQTL (purple).

**Figure S9:**
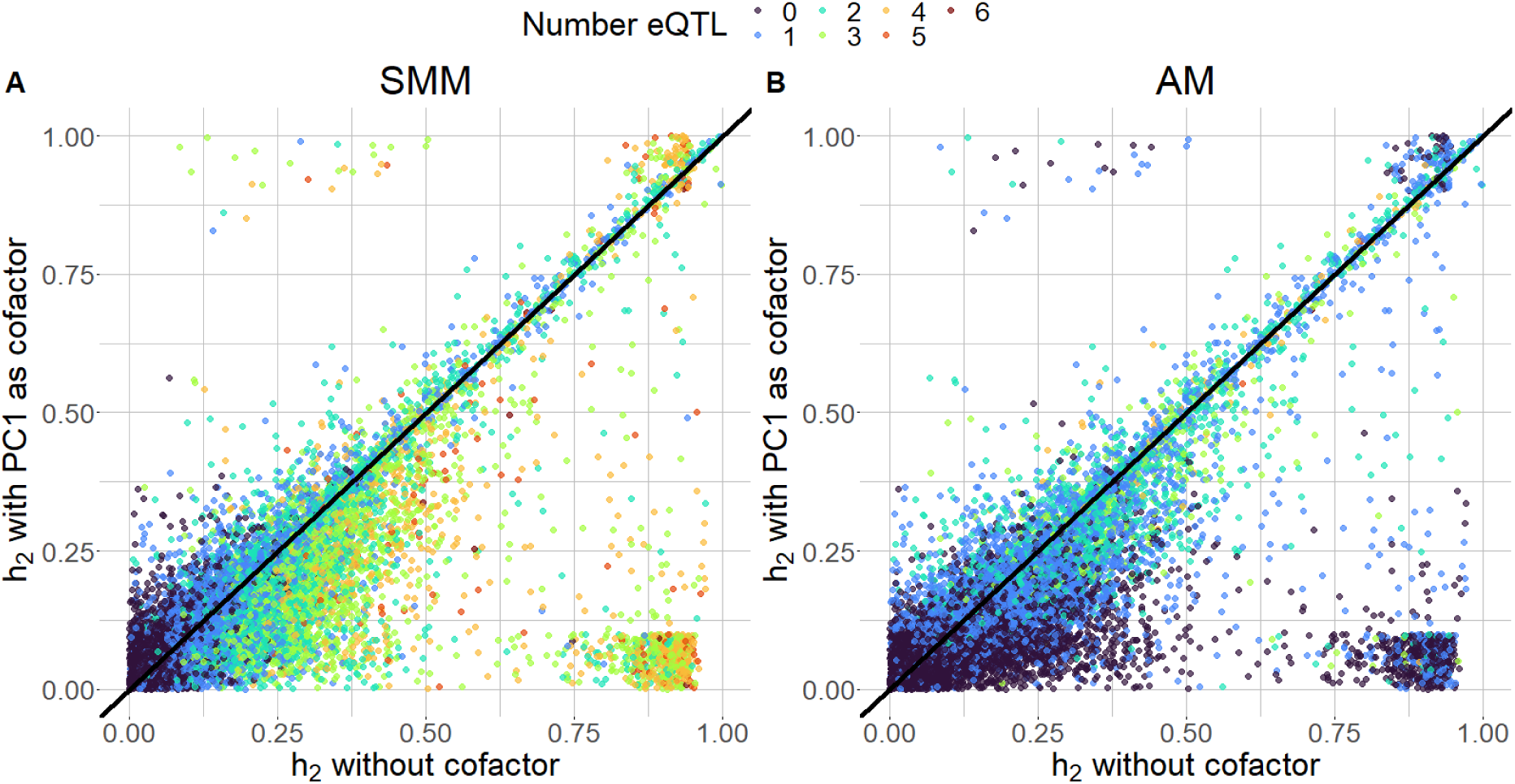
Relationship between heritability of gene expression with and without PC1 as cofactor and the number of eQTLs with the SMM and AAM. The x-axis shows the heritability calculated without PC1 as covariate. The y-axis shows the heritability calculated with PC1 as covariate. Each dot represents one of the 12029 transcripts with expression in our dataset. Black line has slope 1 and passes through the origin. Transcripts are colored by the number of eQTL affecting their expression. Values range from 0 to 6 because we call a maximum of one eQTL per chromosome. A) Number of eQTLs detected with SMM. B) Number of eQTLs detected with AAM.

**Figure S10:**
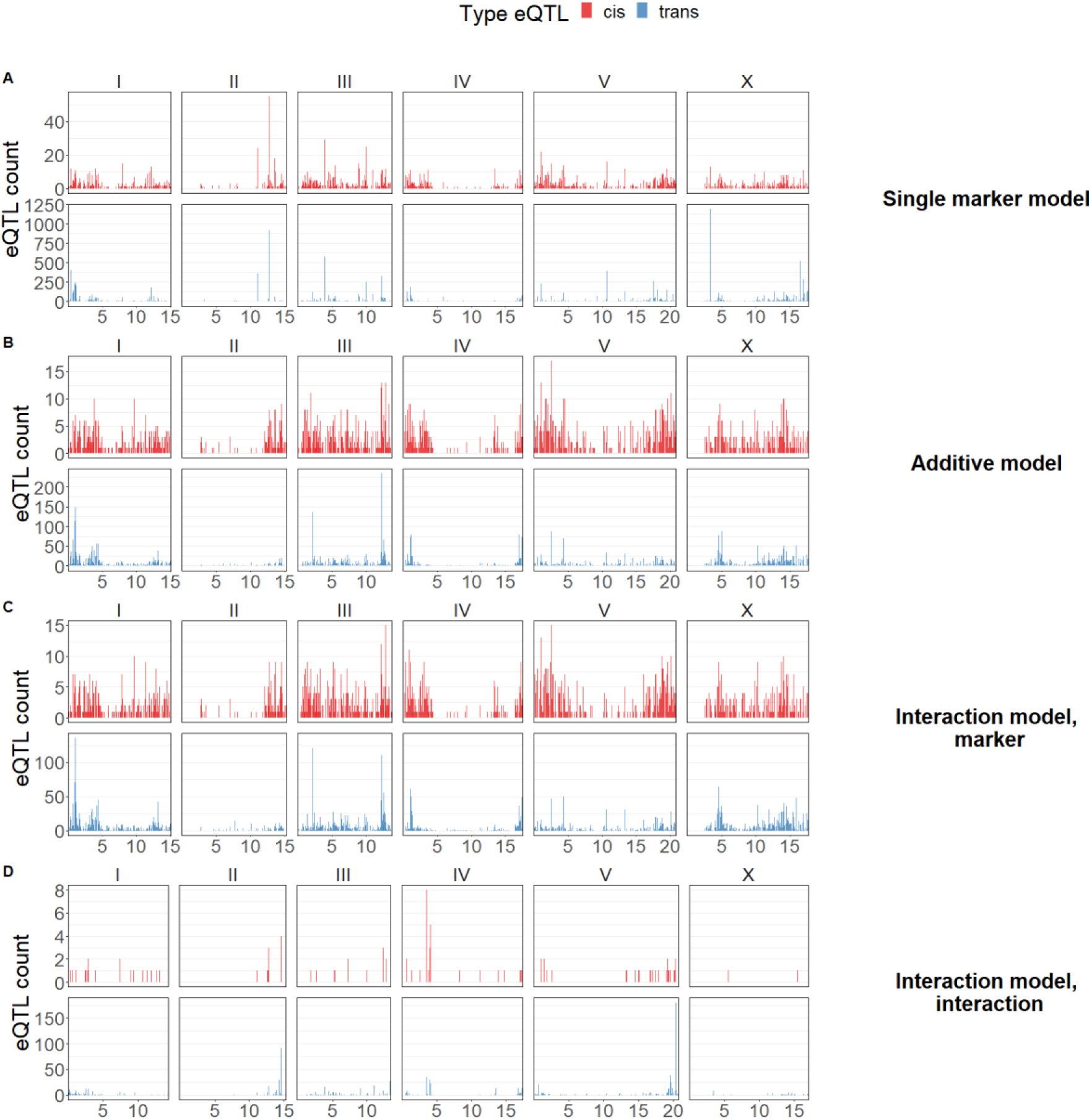
C*i*s/*trans* distribution of eQTLs over the genome. Blue and red colors indicate *trans-e*QTLs and *cis*-eQTLs respectively. Note that the range of the y-axis differs between and within the sub figures.

**Figure S11:**
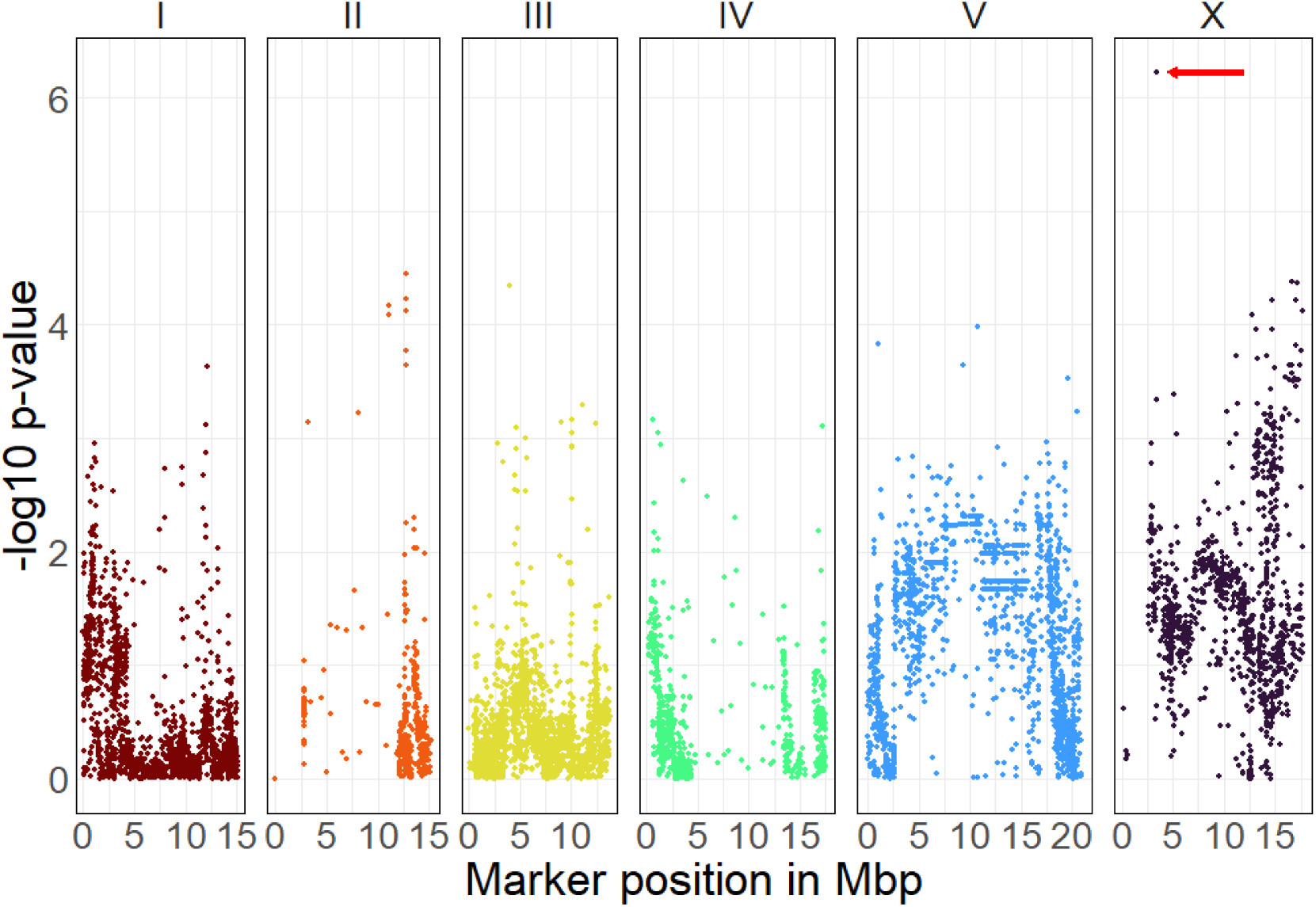
Marker p-values obtained with a linear model of PC1 ∼ marker genotype. The red arrow denotes an outlier corresponding to the marker at 3.404162 Mb on chromosome X. This is also the position of the largest hotspot according to the SMM.

**Figure S12:**
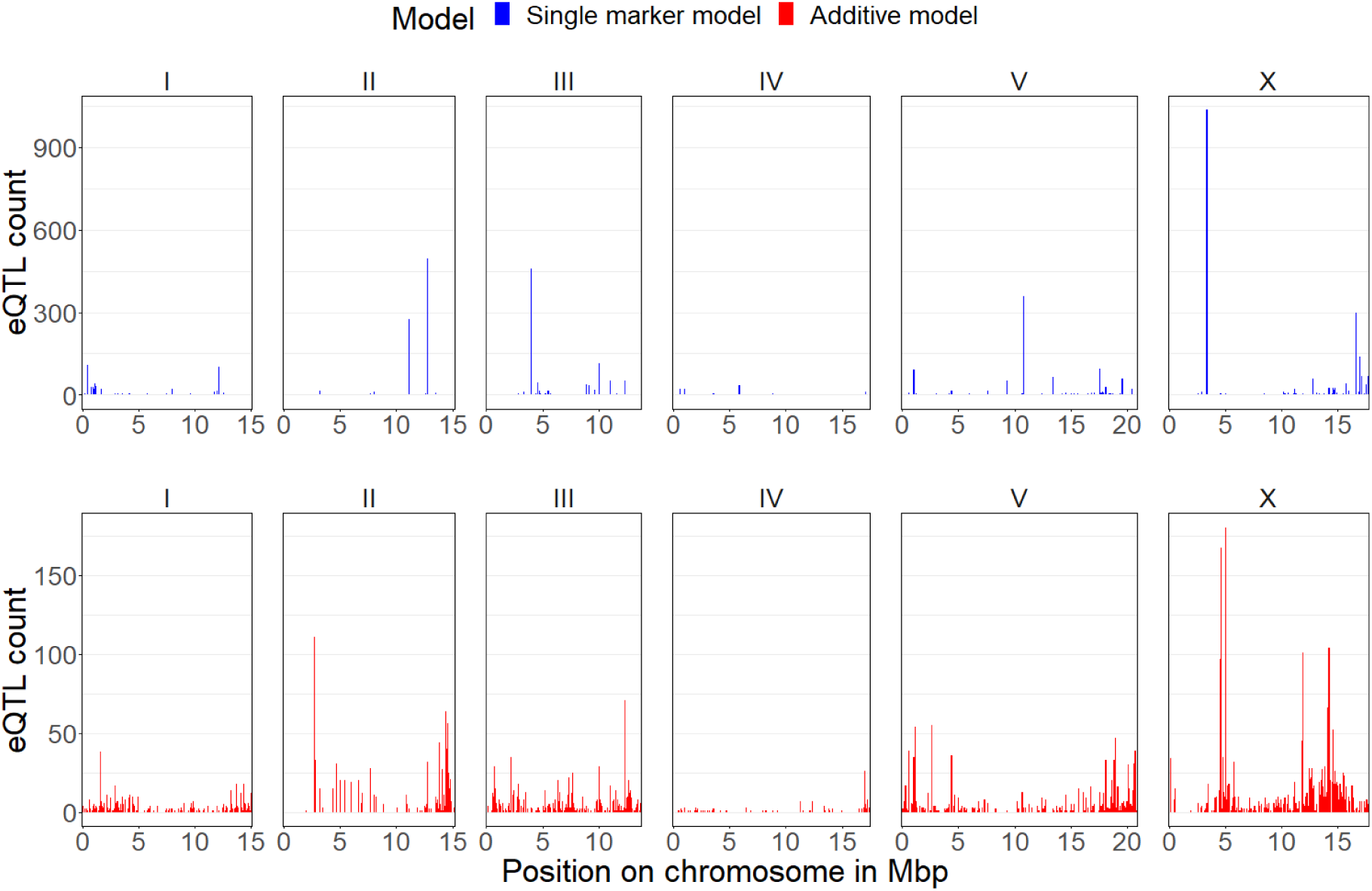
Distribution of single marker eQTLs over the genome. Top panel shows markers with lowest p-value according to SMM (top panel). Bottom panel shows markers with lowest p-value on the same chromosome for the same transcripts according to AAM. Plot only includes eQTLs where the location of the marker with the lowest p-value changes more than 1MB between the SMM and AAM. Note that, by definition of single marker eQTLs, the top panel represents only significant eQTLs, whereas none of the markers in the bottom panel are significantly associated with gene expression.

**Figure S13:**
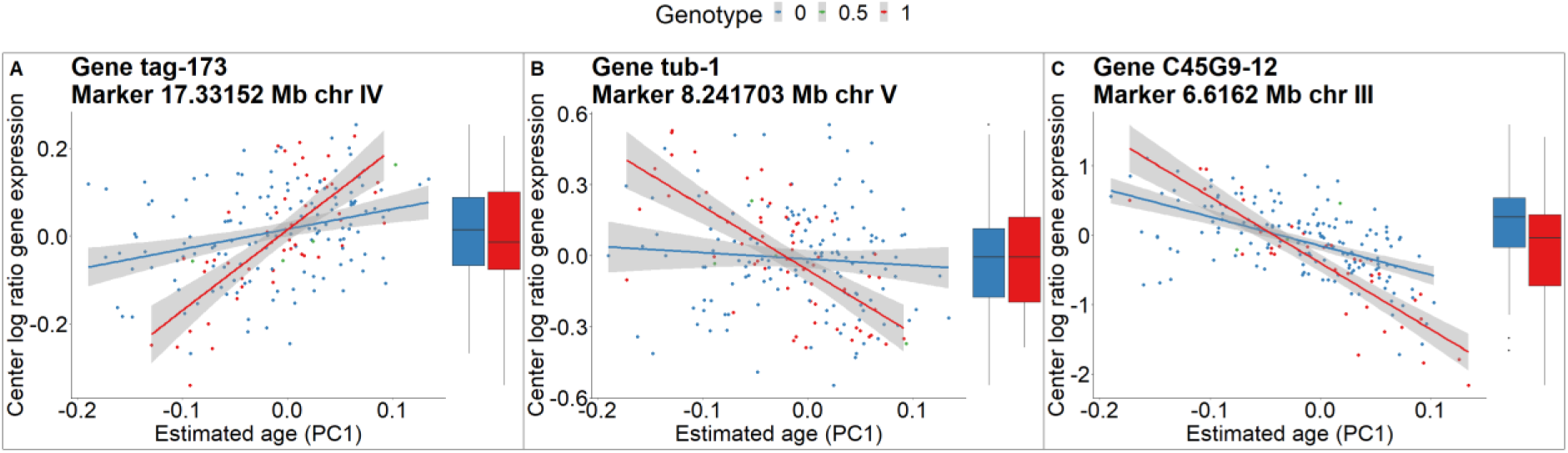
Examples of eQTLs with an interaction that is significant at 0.1 FDR but not at 0.05 FDR. Color corresponds to genotype. Lines show the best linear fit to the data. Shaded areas show the 95% confidence interval. Boxplots show magnitude of gene expression without developmental context.

**Figure S14:**
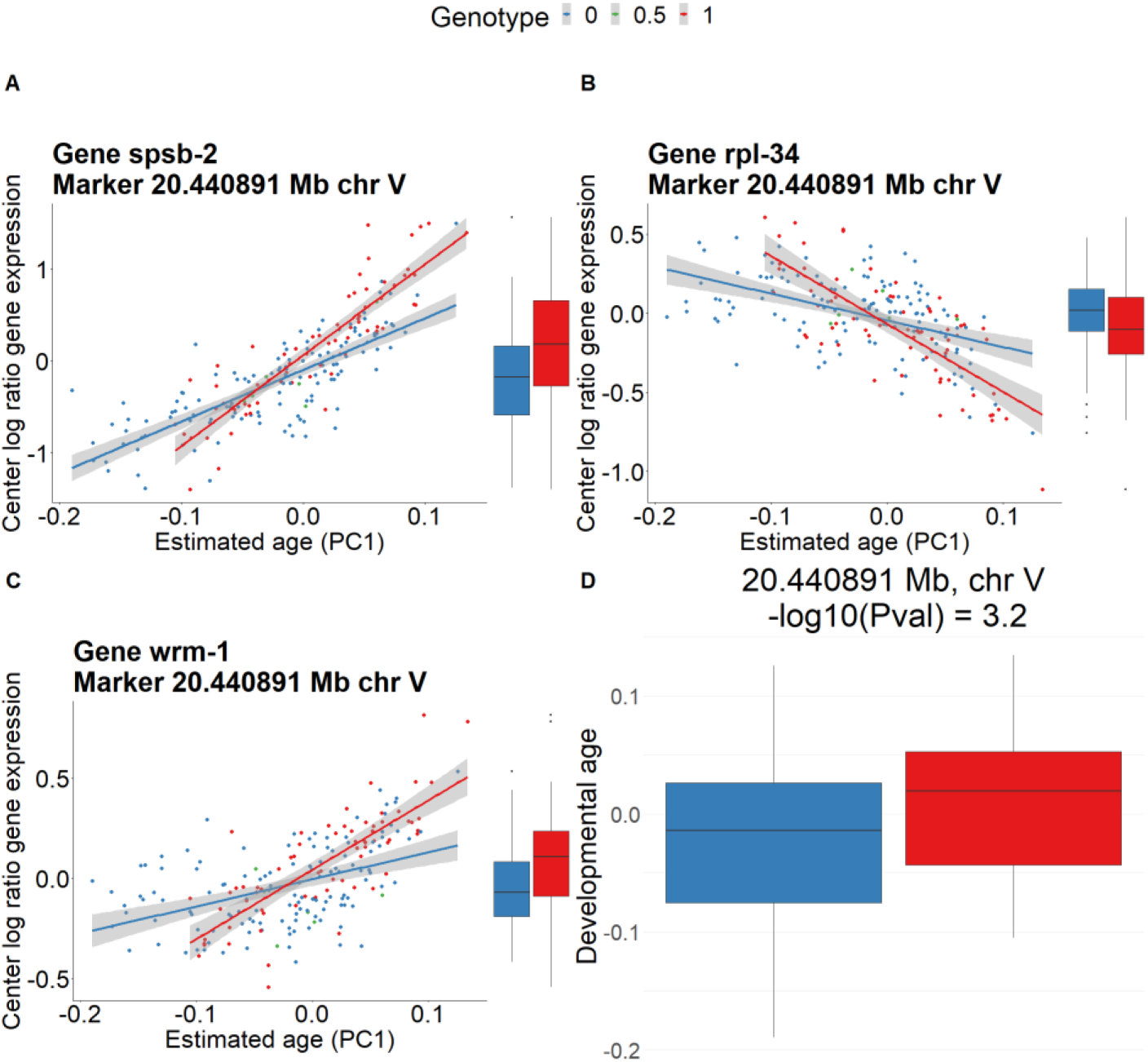
Interaction hotspot on chromosome V affects the rate at which transcript levels change. Hotspot location is at 20.440891 Mb. **A-C)** Expression of three example transcripts with a significant interaction eQTL that maps to the hotspot position. **D)** Boxplot of the developmental age for the alleles at the hotspot marker. The weight of the distribution is skewed towards lower developmental ages for allele 0. P-value is obtained using a linear model of PC1∼genotype at hotspot position.

**Figure S15:**
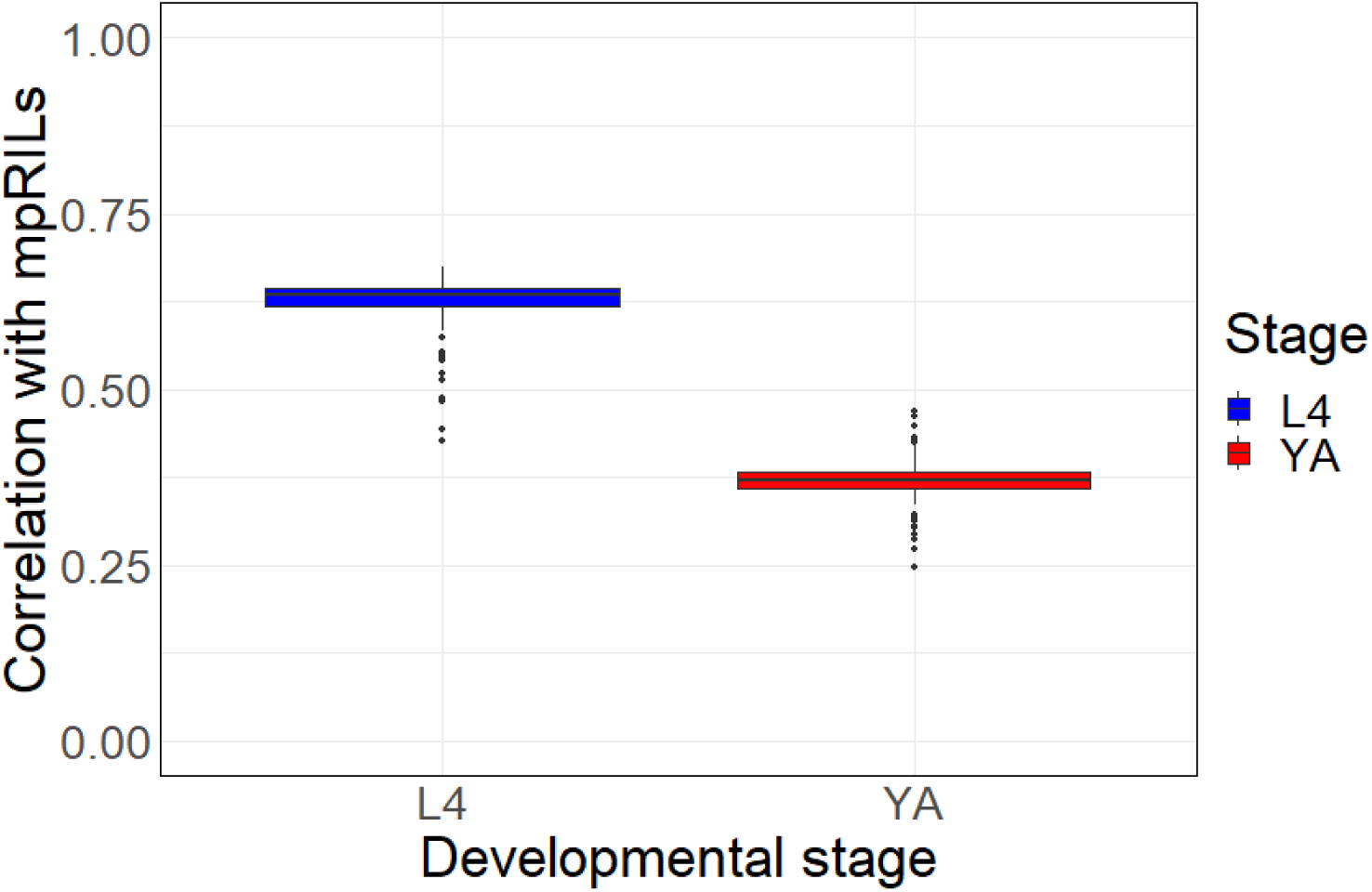
Correlation transcriptomes mpRILs with L4 and young adult reference transcriptomes. Boxplot of the mean correlation of the mpRILs with two L4 reference genomes (L4 and L4b from Boeck *et al.*, 2016) (Blue), and the mean correlation with two young adult reference genomes (YA and N2Yad-1 from Boeck *et al.*, 2016) (Red). For each mpRIL the transcriptome shows a stronger mean correlation with the L4 reference transcriptome compared to the correlation with the L3 reference transcriptome.

**Figure S16:**
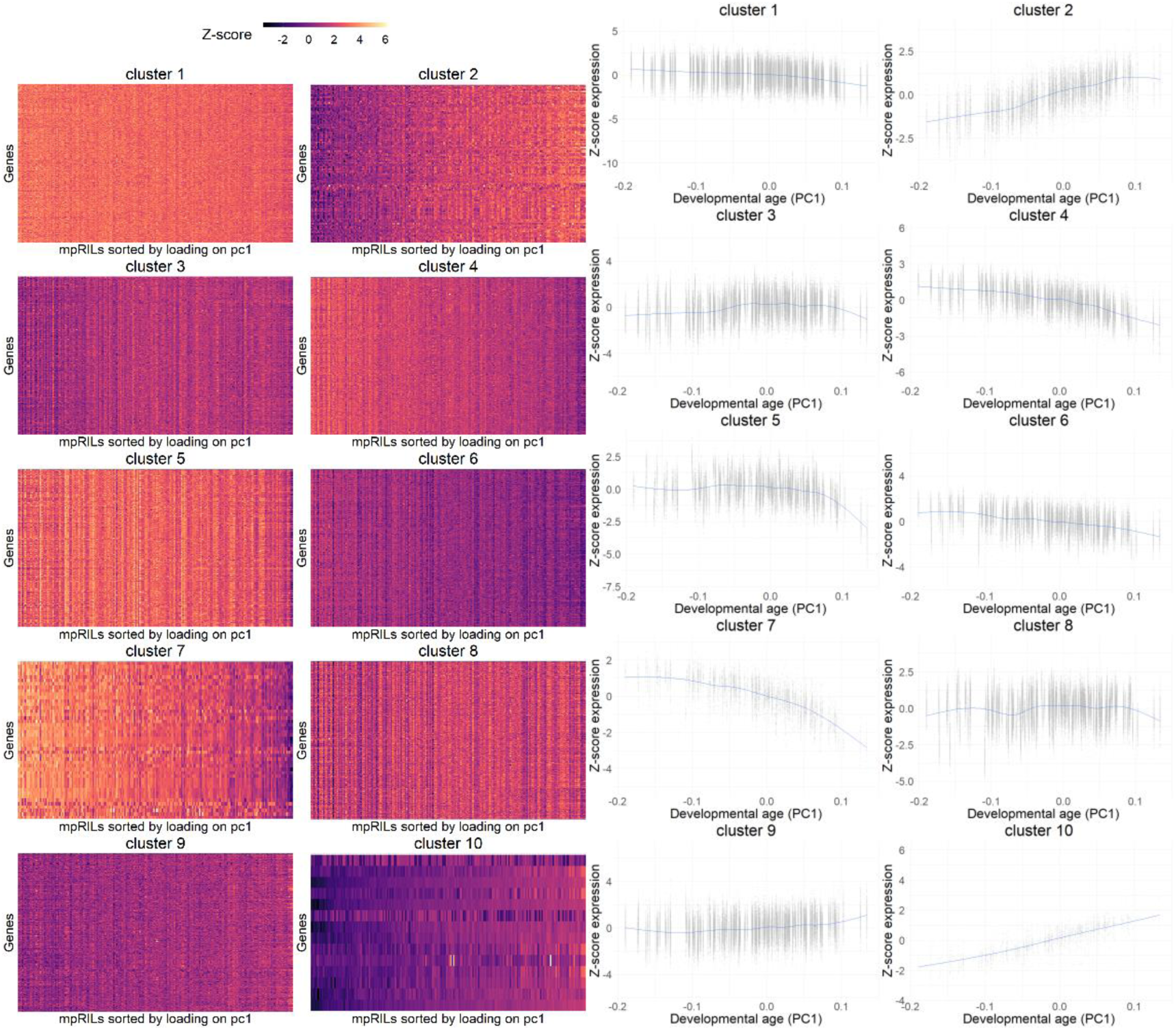
Heatmaps and line plots of Z-scores of center log ratio of gene expression for clusters of oscillatory genes^20^. Clusters were determined using k-means clustering (n = 10) on a gene expression matrix of oscillatory genes. In total 2346 oscillatory genes with significant expression in our data were used in the clustering. The number of genes per cluster from cluster 1 to cluster 10 were 562, 124, 351, 266, 191, 215, 46, 304, 273 and 14.

**Figure S17:**
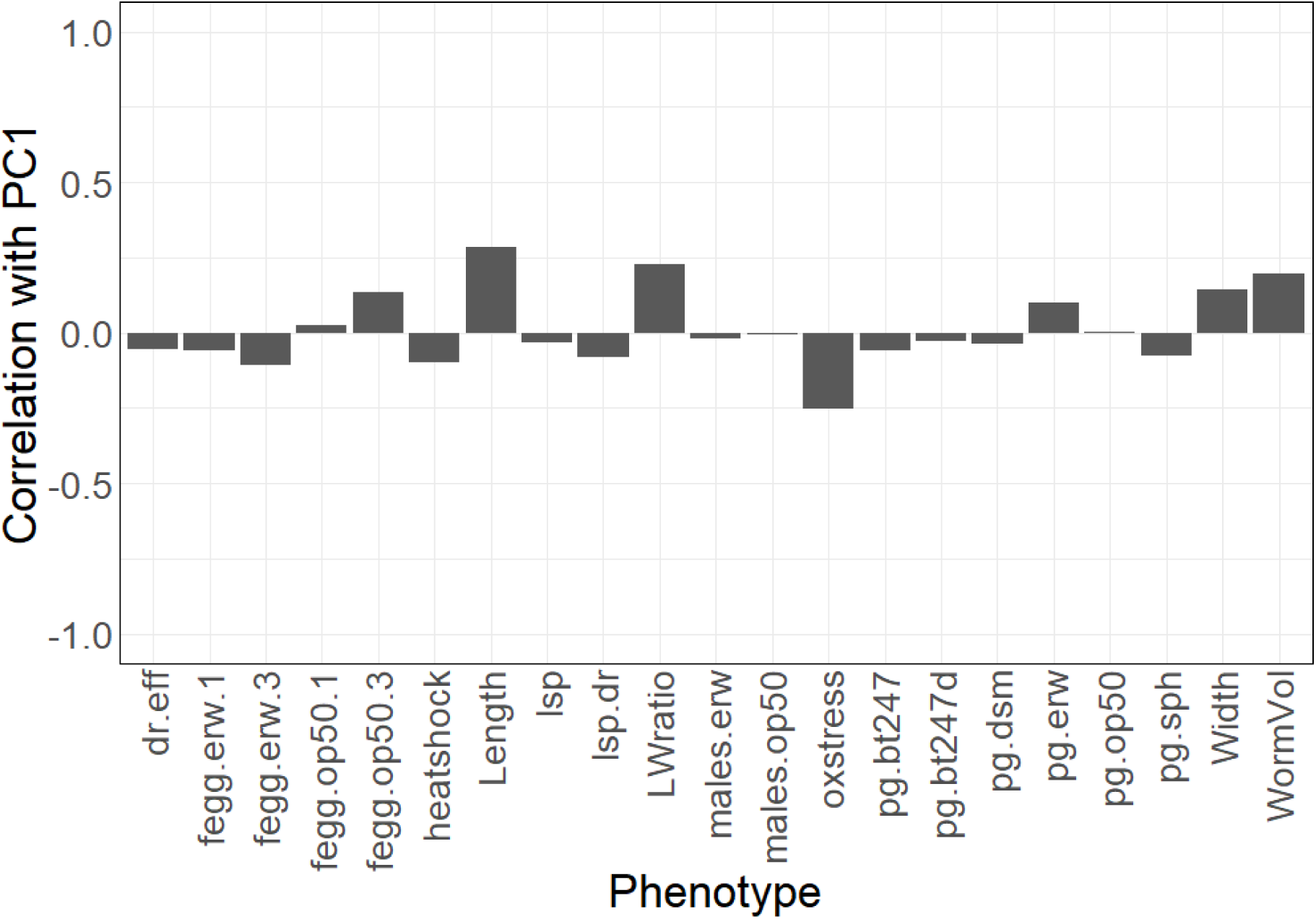
Correlation between phenotypes scored in Snoek *et al.*, 2019 and PC1. From the method section of Snoek *et al.*, 2019: “Dr.eff is the difference in average life span between NGM and DR medium in days. Fegg.erw.1 is the time in hours until the first egg (1–10) for populations grown on Erwinia. Fegg.erw.3 is the time in hours until the first egg (> 100) for populations grown on Erwinia. Fegg.op50.1 is the time in hours until the first egg (1–10) for populations grown on OP50. Fegg.op50.3 is the time in hours until the first egg (> 100) for populations grown on OP50. Heat shock is the average number of dead animals per 50. Length is in nanometers. Lsp is the average lifespan on NGM in days. lsp.dr is the average lifespan on DR medium in days. LWratio is the length in nanometers divided by the width in nanometers. Oxidative stress indicates activity in terms of movement after addition of hydrogen peroxide. Males.erw and males.op50 is the occurrence of males on plates (0 = none, 0.5 = 1 plate, 1 = 2 plates). Pg.bt247, pgbt247d, pg.dsm, pg.erw, pg.op50 and pg.sph shows population growth (worms per 5 μl of culture) on a pathogenic Bacillus thuringiensis strain NRRL B-18247 on two concentrations of 1:300 and 1:600, a non-pathogenic Bacillus thuringiensis strain DSM-350E, Erwinia rhapontici (isolated from Orsay, France), Escherichia coli OP50 and Sphingobacterium sp. (isolated from Orsay, France). Width is in nanometers. Wormvolume is volume in nanoliters”.

## Supplementary text A1: Investigating allele dependent non-linear gene expression dynamics using a natural spline model

### Methods

To perform eQTL mapping with the natural splines we used the ns() function from the splines package to model the developmental age. We used three knots, chosen such that each interval characterized by the knots contains one fourth of the mpRILs. We obtained p-values by performing a two model anova using the anova() function (stats package, base R). One of the models included in the anova contained an interaction between the marker and a natural spline of the developmental age while the other contained only an additive contribution of the natural spline. We used a cutoff of –log10(P) = 7.53 resulting in an FDR of 0.2, as using an FDR of 0.1 (-log10(P) = 12.1) only resulted in 25 significant interactions, while it was impossible to threshold at an FDR of 0.05.

### Results

Gene expression dynamics can be more complex than simple linear trajectories. To investigate the prevalence of allele dependent non-linear relationships between gene expression and PC1 we performed eQTL mapping using a natural spline of the developmental age^2^. To search for non-linear interactions between the developmental age and the genotype we used a cutoff of –log10(P) > 7.53 (FDR = 0.2). More stringent cutoffs detect very few interactions. The 0.2 FDR threshold revealed the presence of 113 interesting non-linear gene expression patterns **(Figure A1.1**). Furthermore, we found multiple qualitatively similar expression patterns in transcripts associated with the same dynamical eQTL **(Figure A1.2**), demonstrating that complex dynamics can be heritable.

To quantify the effect of eQTL detection with the natural spline model we divided the 113 eQTLs into categories **(Figure A1.3**). The first category (55 eQTLs) contains those eQTLs which clearly result from highly uneven genotype distributions or just a few outliers. Uneven distributions or outliers can lead to overfitting because of the flexibility of the natural spline model. Furthermore, due to the multi-parental origin of the mpRILs, uneven allelic distributions are prevalent in our data. The second category (25 eQTLs) corresponds to interactions that were approximately linear. Such gene expression patterns are also adequately described by a linear interaction model. A third category (11 eQTLs) contains eQTLs with a non-linear but approximately monotonic gene expression pattern. Finally, the fourth category (22 eQTLs) consists of non-monotonic gene expression patterns. Especially eQTLs in this latter category demonstrate the added value of natural splines, as complex patterns can average out to a similar magnitude (**Figure A1.1C**).

**Figure A1.1:**
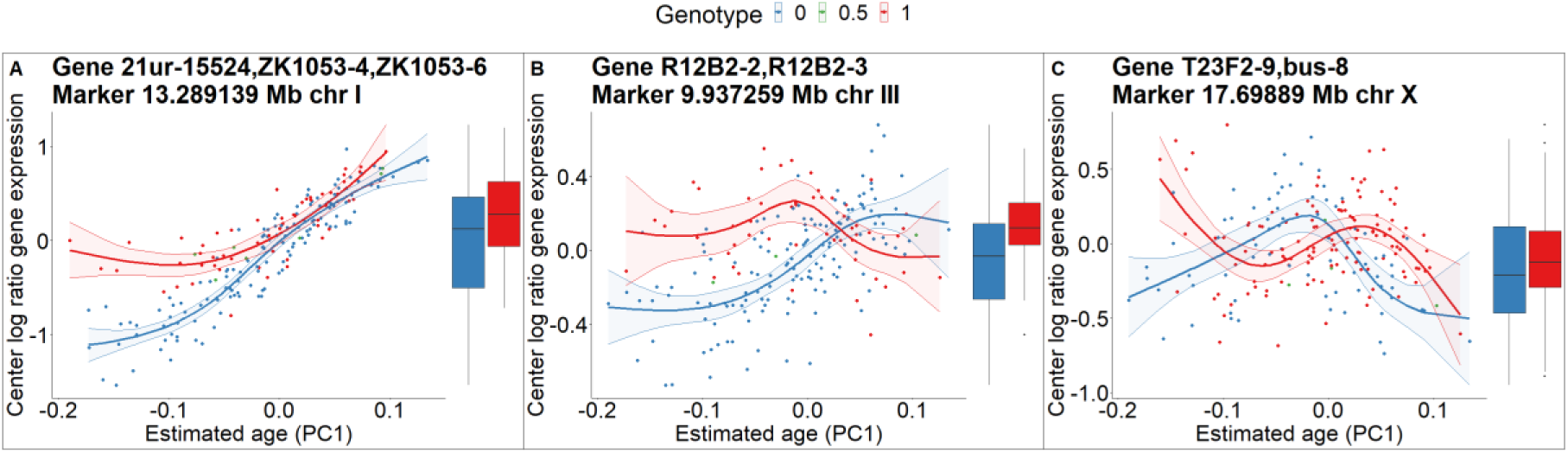
Example non-linear gene expression patterns detected using natural splines. Colors correspond to genotype at eQTL position. Lines are best fit of natural spline model (shaded area is 95% CI) and show the dynamics of gene expression. Boxplots show magnitude of gene expression without temporal component. **A)** eQTL that causes disparate monotonic expression dynamics at the lower end of the developmental age range of the mpRILs. **B)** eQTL for which one genotype shows a monotonic increase over the developmental age while the other shows non-monotonic concave dynamics. **C)** eQTL where both genotypes show different complex, non-monotonic dynamics.

**Figure A1.2:**
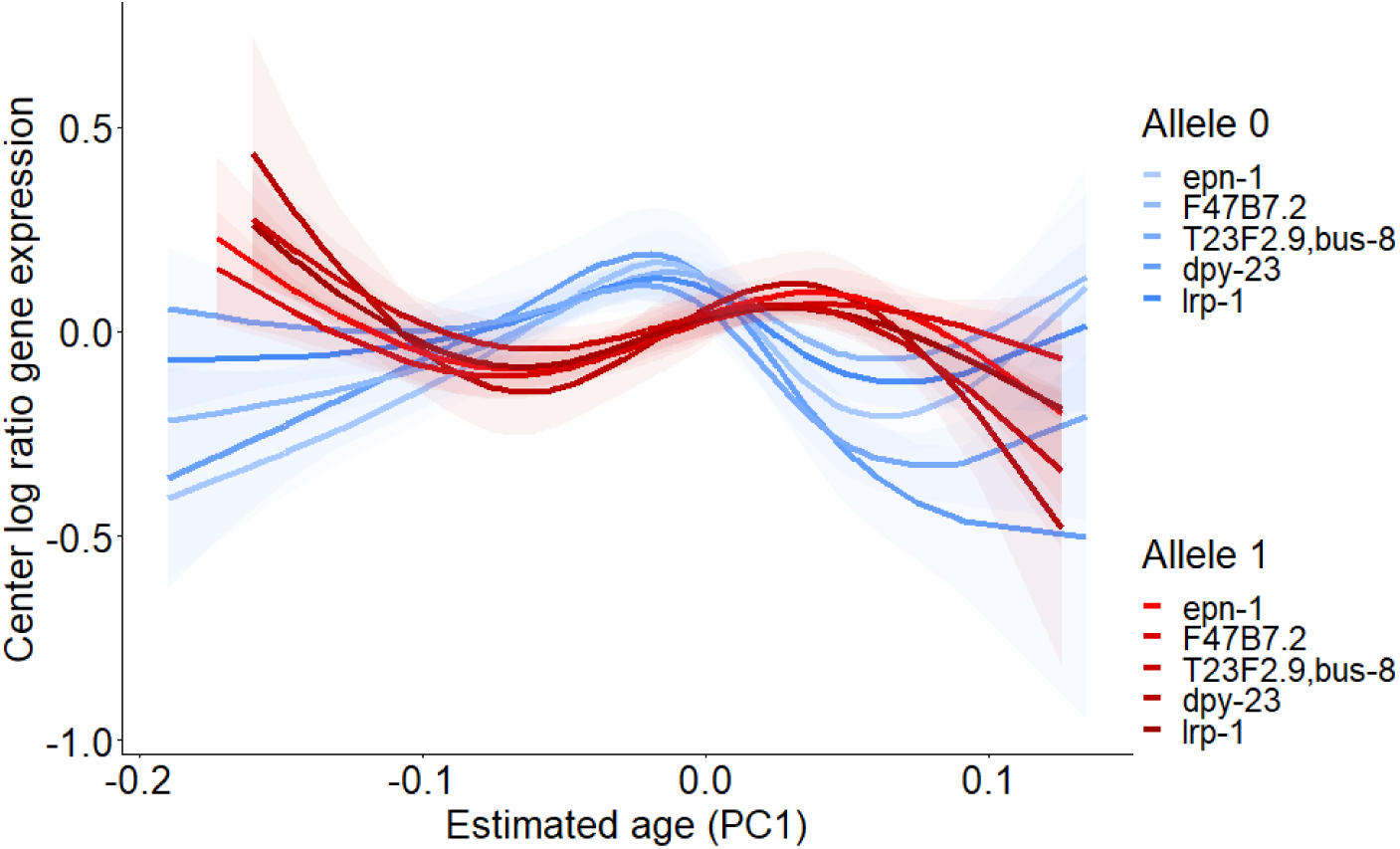
Example of marker causing consistent dynamics for multiple transcripts. Fit of the natural spline model to the center log ratio of gene expression of transcripts with a –log10(interaction p-value) >7.5 in the spline models with the adjacent markers at 17.519998 Mb chr X (*F47B7.2*; *lrp-1*) or 17.69889 Mb chr X (*epn- 1*; *T23F2.9*, *bus−8*; *dpy-23*).

**Figure A1.3:**
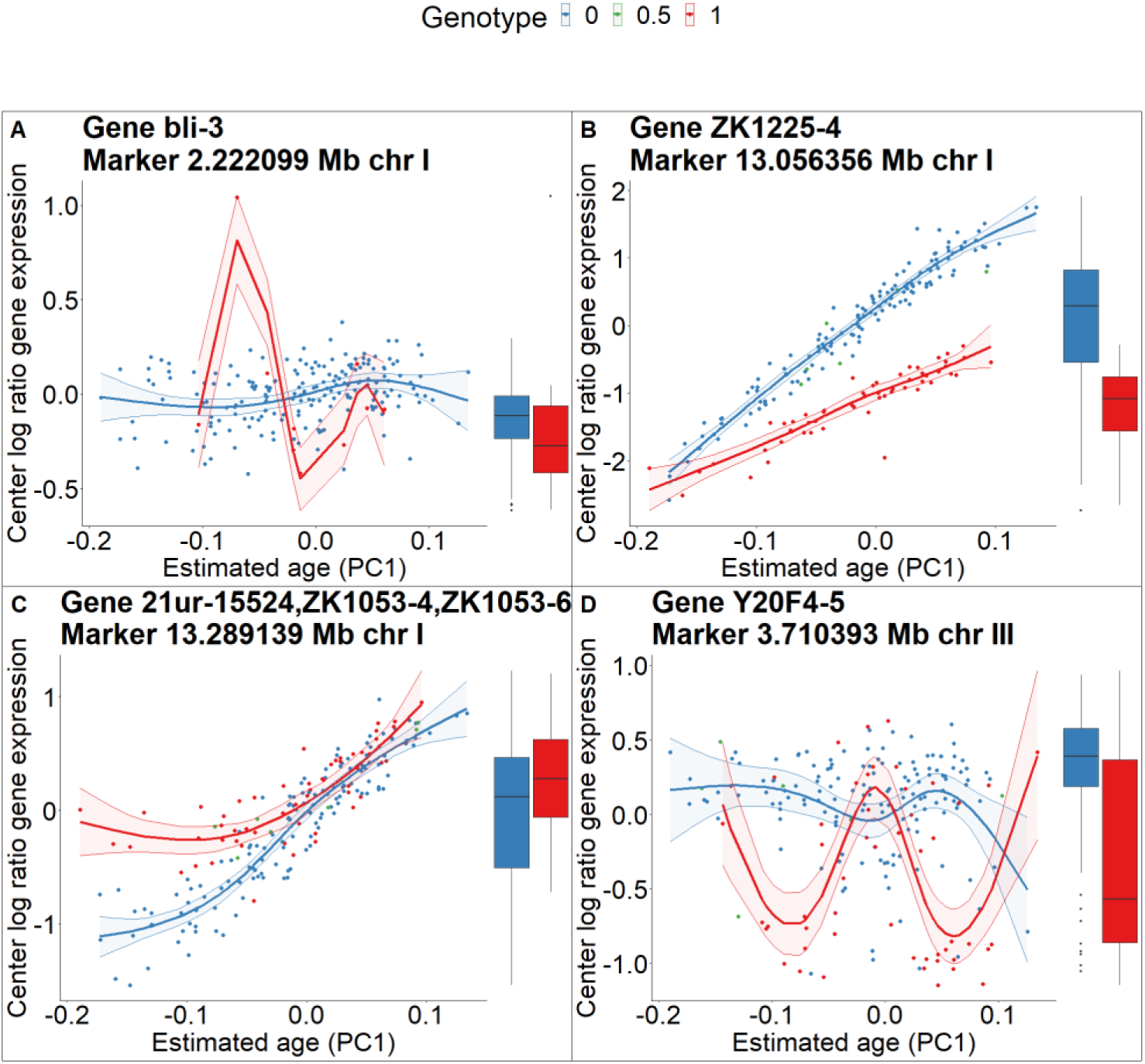
Four categories of eQTLs detected using the interaction term of the natural spline model at 0.2 FDR. Category was scored by eye from a total of 113 eQTLs. **A)** False positive due to uneven genotype distribution/outlier. **B)** Approximately linear interaction. **C)** Monotonic non-linear interaction. **D)** Non-monotonic interaction. We excluded from the scoring two significant markers for which almost all mpRILs were heterozygous.

## References

1. Snoek, L. B., Sterken, M. G., Volkers, R. J. M., Klatter, M., Bosman, K. J., Bevers, R. P. J., Riksen, J. A. G., Smant, G., Cossins, A. R. & Kammenga, J. E. A rapid and massive gene expression shift marking adolescent transition in C. elegans. Scientific Reports 2014 4:1 4, 1–5 (2014).

2. Francesconi, M. & Lehner, B. The effects of genetic variation on gene expression dynamics during development. Nature 2014 505:7482 505, 208–211 (2014).

3. Jovic, K., Sterken, M. G., Grilli, J., Bevers, R. P. J., Rodriguez, M., Riksen, J. A. G., Allesina, S., Kammenga, J. E. & Snoek, L. B. Temporal dynamics of gene expression in heat-stressed Caenorhabditis elegans. PLoS One 12, e0189445 (2017).

4. Spencer, W. C. et al. A spatial and temporal map of C. elegans gene expression. Genome Res 21, (2011).

5. Araya, C. L. et al. Regulatory analysis of the C. Elegans genome with spatiotemporal resolution. Nature 512, (2014).

6. Boeck, M. E., Huynh, C., Gevirtzman, L., Thompson, O. A., Wang, G., Kasper, D. M., Reinke, V., Hillier, L. W. & Waterston, R. H. The time-resolved transcriptome of C. Elegans. Genome Res 26, 1441–1450 (2016).

7. Reinke, V., Smith, H. E., Nance, J., Wang, J., Van Doren, C., Begley, R., Jones, S. J. M., Davis, E. B., Scherer, S., Ward, S. & Kim, S. K. A Global Profile of Germline Gene Expression in C. elegans. Mol Cell 6, 605–616 (2000).

8. Jansen, R. C. & Nap, J. P. Genetical genomics: the added value from segregation. Trends in Genetics 17, 388–391 (2001).

9. Snoek, L. B., Van Der Velde, K. J., Arends, D., Li, Y., Beyer, A., Elvin, M., Fisher, J., Hajnal, A., Hengartner, M. O., Poulin, G. B., Rodriguez, M., Schmid, T., Schrimpf, S., Xue, F., Jansen, R. C., Kammenga, J. E. & Swertz, M. A. WormQTL—public archive and analysis web portal for natural variation data in Caenorhabditis spp. Nucleic Acids Res 41, D738 (2013).

10. Snoek, B. L., Sterken, M. G., Hartanto, M., Van Zuilichem, A. J., Kammenga, J. E., De Ridder, D. & Nijveen, H. WormQTL2: an interactive platform for systems genetics in Caenorhabditis elegans. Database (Oxford) 2020, 149 (2020).

11. Evans, K. S., van Wijk, M. H., McGrath, P. T., Andersen, E. C. & Sterken, M. G. From QTL to gene: C. elegans facilitates discoveries of the genetic mechanisms underlying natural variation. Trends Genet 37, 933 (2021).

12. Andersen, E. C. & Rockman, M. v. Natural genetic variation as a tool for discovery in Caenorhabditis nematodes. Genetics 220, (2022).

13. Sterken, M. G., Snoek, L. B., Kammenga, J. E. & Andersen, E. C. The laboratory domestication of Caenorhabditis elegans. Trends in Genetics 31, 224–231 (2015).

14. Gaertner, B. E. & Phillips, P. C. Caenorhabditis elegans as a platform for molecular quantitative genetics and the systems biology of natural variation. Center for Ecology and Evolutionary Biology (2010) doi:10.1017/S0016672310000601.

15. Hashimshony, T., Feder, M., Levin, M., Hall, B. K. & Yanai, I. Spatiotemporal transcriptomics reveals the evolutionary history of the endoderm germ layer. Nature vol. 519 Preprint at https://doi.org/10.1038/nature13996 (2015).

16. Levin, M. et al. The mid-developmental transition and the evolution of animal body plans. Nature 531, 637–641 (2016).

17. McCarroll, S. A., Murphy, C. T., Zou, S., Pletcher, S. D., Chin, C.-S., Nung Jan, Y., Kenyon, C., Bargmann, C. I. & Li, H. Comparing genomic expression patterns across species identifies shared transcriptional profile in aging. Nat Genet 9, (2004).

18. Viñuela, A., Snoek, L. B., Riksen, J. A. G. & Kammenga, J. E. Genome-wide gene expression regulation as a function of genotype and age in C. elegans. Genome Res 20, 929–937 (2010).

19. Viñuela, A., Basten Snoek, L., Riksen, J. A. G. & Kammenga, J. E. Aging uncouples heritability and expression-qtl in caenorhabditis elegans. G3: Genes, Genomes, Genetics 2, 597–605 (2012).

20. Hendriks, G. J., Gaidatzis, D., Aeschimann, F. & Großhans, H. Extensive Oscillatory Gene Expression during C. elegans Larval Development. Mol Cell 53, 380–392 (2014).

21. Meeuse, M. W., Hauser, Y. P., Morales Moya, L. J., Hendriks, G., Eglinger, J., Bogaarts, G., Tsiairis, C. & Großhans, H. Developmental function and state transitions of a gene expression oscillator in Caenorhabditis elegans . Mol Syst Biol 16, (2020).

22. Kim, D. H., Grün, D. & van Oudenaarden, A. Dampening of expression oscillations by synchronous regulation of a microRNA and its target. Nat Genet 45, (2013).

23. Bulteau, R. & Francesconi, M. Real age prediction from the transcriptome with RAPToR. Nature Methods 2022 19:8 19, 969–975 (2022).

24. Snoek, B. L., Volkers, R. J. M., Nijveen, H., Petersen, C., Dirksen, P., Sterken, M. G., Nakad, R., Riksen, J. A. G., Rosenstiel, P., Stastna, J. J., Braeckman, B. P., Harvey, S. C., Schulenburg, H. & Kammenga, J. E. A multi-parent recombinant inbred line population of C. elegans allows identification of novel QTLs for complex life history traits. BMC Biol 17, (2019).

25. Volkers, R. J. M., Snoek, L. B., Hubar, C. J. van H., Coopman, R., Chen, W., Yang, W., Sterken, M. G., Schulenburg, H., Braeckman, B. P. & Kammenga, J. E. Gene-environment and protein- degradation signatures characterize genomic and phenotypic diversity in wild Caenorhabditis elegans populations. BMC Biol 11, 1–13 (2013).

26. Perez, M. F., Francesconi, M., Hidalgo-Carcedo, C. & Lehner, B. Maternal age generates phenotypic variation in Caenorhabditis elegans. Nature 552, (2017).

27. Ben-David, E., Boocock, J., Guo, L., Zdraljevic, S., Bloom, J. S. & Kruglyak, L. Whole-organism eqtl mapping at cellular resolution with single-cell sequencing. Elife 10, (2021).

28. Jovic, K., Grilli, J., Sterken, M. G., Snoek, B. L., Riksen, J. A. G., Allesina, S. & Kammenga, J. E. Transcriptome resilience predicts thermotolerance in Caenorhabditis elegans. BMC Biol 17, 1–12 (2019).

29. Snoek, B. L., Sterken, M. G., Nijveen, H., Volkers, R. J. M., Riksen, J., Rosenstiel, P. C., Schulenburg, H. & Kammenga, J. E. The genetics of gene expression in a Caenorhabditis elegans multiparental recombinant inbred line population. G3 Genes|Genomes|Genetics 11, (2021).

30. Kamath, R. S., Fraser, A. G., Dong, Y., Poulin, G., Durbin, R., Gotta, M., Kanapin, A., le Bot, N., Moreno, S., Sohrmann, M., Welchman, D. P., Zipperien, P. & Ahringer, J. Systematic functional analysis of the Caenorhabditis elegans genome using RNAi. Nature 2003 421:6920 421, 231–237 (2003).

31. Kenyon, C., Chang, J., Gensch, E., Rudner, A. & Tabtiang, R. A C. elegans mutant that lives twice as long as wild type. Nature 366, (1993).

32. Breitling, R., Li, Y., Tesson, B. M., Fu, J., Wu, C., Wiltshire, T., Gerrits, A., Bystrykh, L. v., de Haan, G., Su, A. I. & Jansen, R. C. Genetical genomics: Spotlight on QTL hotspots. PLoS Genetics vol. 4 Preprint at https://doi.org/10.1371/journal.pgen.1000232> (2008).

33. Michaelson, J. J., Loguercio, S. & Beyer, A. Detection and interpretation of expression quantitative trait loci (eQTL). Methods vol. 48 Preprint at https://doi.org/10.1016/j.ymeth.2009.03.004 (2009).

34. Mata-Cabana, A., Romero-Expósito, F. J., Geibel, M., Piubeli, F. A., Merrow, M. & Olmedo, M. Deviations from temporal scaling support a stage-specific regulation for C. elegans postembryonic development. BMC Biol 20, 94 (2022).

35. Sterken, M. G., Bevers, R. P. J., Volkers, R. J. M., Riksen, J. A. G., Kammenga, J. E. & Snoek, B. L. Dissecting the eQTL Micro-Architecture in Caenorhabditis elegans. Front Genet 11, (2020).

36. Snoek, B. L., Sterken, M. G., Bevers, R. P. J., Volkers, R. J. M., van’t Hof, A., Brenchley, R., Riksen, J. A. G., Cossins, A. & Kammenga, J. E. Contribution of trans regulatory eQTL to cryptic genetic variation in C. elegans. BMC Genomics 18, (2017).

37. Li, Y., Álvarez, O. A., Gutteling, E. W., Tijsterman, M., Fu, J., Riksen, J. A. G., Hazendonk, E., Prins, P., Plasterk, R. H. A., Jansen, R. C., Breitling, R. & Kammenga, J. E. Mapping determinants of gene expression plasticity by genetical genomics in C. elegans. PLoS Genet 2, (2006).

38. Li, Y., Breitling, R., Snoek, L. B., Van Der Velde, K. J., Swertz, M. A., Riksen, J., Jansen, R. C. & Kammenga, J. E. Global genetic robustness of the alternative splicing machinery in Caenorhabditis elegans. Genetics 186, (2010).

39. Sterken, M. G., van der Plaat, L. van B., Riksen, J. A. G., Rodriguez, M., Schmid, T., Hajnal, A., Kammenga, J. E. & Snoek, B. L. Ras/MAPK modifier loci revealed by eQTL in Caenorhabditis elegans. G3: Genes, Genomes, Genetics 7, (2017).

40. van Wijk, M. H., Riksen, J. A. G., Elvin, M., Poulin, G. B., Maulana, M. I., Kammenga, J. E., Snoek, B. L. & Sterken, M. G. Cryptic genetic variation of eQTL architecture revealed by genetic perturbation in C. elegans. G3 Genes|Genomes|Genetics (2023) doi:10.1093/g3journal/jkad050.

41. Filina, O., Demirbas, B., Haagmans, R. & van Zon, J. S. Temporal scaling in C. elegans larval development. Proc Natl Acad Sci U S A 119, (2022).

42. O’Duibhir, E., Lijnzaad, P., Benschop, J. J., Lenstra, T. L., Leenen, D., Groot Koerkamp, M. J., Margaritis, T., Brok, M. O., Kemmeren, P. & Holstege, F. C. Cell cycle population effects in perturbation studies. Mol Syst Biol 10, (2014).

43. R Core Team (2022). R: A language and environment for statistical computing. R Foundation for Statistical Computing. Preprint at (2022).

44. Martin, V., Schbath, S. & Hennequet-Antier, C. R graphics with ggplot2. (2022).

